# Megalin-mediated deterministic pseudotransduction in proximal tubules fundamentally challenges current understanding of AAV kidney transduction

**DOI:** 10.64898/2026.10.26.761697

**Authors:** Anusha Sairavi, Ranjan Das, Lena Li, Tahnee Groat, Yasunori Matsuzaki, Hirokazu Hirai, Michael Hutchens, Hiroyuki Nakai

## Abstract

Genetic kidney diseases are promising targets for gene therapy. Emerging AAV capsids that reportedly enable effective gene delivery to proximal tubules (PTs) following intravenous injection have raised hopes for treating diseases affecting renal tubules. Despite these promising developments, the scientific literature contains contradictory and non-reproducible findings, which prompted us to conduct a detailed and rigorous investigation of the mechanisms of AAV vector-mediated PT transduction. Here we provide compelling evidence that previously reported effective PT transduction following intravenous AAV-GFP or AAV-mCherry administration largely reflects megalin-mediated deterministic pseudotransduction rather than stochastic vector-mediated gene delivery. We demonstrate that this process is mediated by megalin-dependent uptake of circulating reporter proteins released from transduced organs including the liver and filtered through the glomeruli. These observations highlight the ongoing challenge of achieving true proximal tubule transduction via systemic AAV delivery and caution against relying on small reporters like GFP or mCherry, which can produce misleading signals.

## Introduction

Gene therapy has transformed the treatment of certain types of genetic diseases in the real-world, exemplified by the success of Zolgensma for spinal muscular atrophy, and continues to show remarkable promise for a wide range of genetic disorders in both preclinical and clinical studies^1^. However, translating gene therapy approaches to genetic kidney diseases has proven particularly challenging. The kidney presents unique challenges for gene delivery due to its complex microanatomy and the presence of specialized physical and functional barriers. In particular, the glomerular filtration barrier, tubular basement membranes, and capillary endothelium around tubules limit the passage of macromolecules, including gene delivery vectors, from the bloodstream and restrict their access to key therapeutic target cells, podocytes and renal tubules. These limitations have slowed progress toward the development of effective renal gene therapies^2, 3, 4, 5^.

Among currently available gene delivery platforms, adeno-associated virus (AAV) vectors have emerged as the most promising for in vivo gene therapy. AAV vectors have established an excellent safety and efficacy record^6, 7, 8^. Since 1999^9^, approximately 40 studies have reported the use of various AAV capsids for kidney-directed gene delivery in rodents, with AAV9 being the most extensively investigated, accounting for more than half of these reports. Although AAV vector-mediated gene delivery has historically been considered inefficient, a careful review of the literature reveals striking study-to-study variabilities, ranging from no detectable to highly efficient transduction, even when the same serotype and administration route were used in the same species. For example, intravenous (IV) injection of AAV9 vectors expressing a reporter gene under the control of a strong ubiquitous promoter resulted in no detectable kidney transduction in mice^10, 11^, whereas other groups reported effective glomerular transduction in the kidney following IV administration of AAV9 vectors at comparable doses^12, 13^.

Reported findings on renal tubular transduction with AAV vectors in mice and non-human primates are further puzzling. We have recently shown that efficient proximal tubule (PT) transduction can be achieved only when transgenes are delivered via the renal pelvis (RP) or renal vein (RV) injection using engineered capsids such as AAV-KP1 and AAV-KP3^14^, but not with AAV9 or other naturally occurring serotypes, or not following IV administration, in healthy mouse kidneys^12, 15^. In addition, we and others have shown that effective PT transduction by IV injection of AAV9 vectors in mice requires impairment of renal physical and functional barriers observed in diseased kidneys such as those with chronic kidney diseases (CKD) in Alport syndrome mice^12, 16^. Others have also shown limited PT transduction with AAV9 in mice following IV, RP or RV injection^13, 17, 18, 19^, conforming to our observations. Despite our own observations and those by others that AAV9 fails to efficiently transduce PTs in mice regardless of the administration route, several groups have reported effective PT transduction with AAV9^20, 21, 22^. There are additional studies that report widespread renal transduction following IV administration of AAV9, which could suggest effective PT transduction, although these studies lack detailed analyses to confirm the specific cell types transduced with AAV9^23, 24^. More recently, engineered capsids such as AAV.cc47^25^, AAV.k13^26^, and AAV.k20^26^, have been reported to mediate robust PT transduction in mice following IV injection^26, 27, 28^. These new capsids have opened a potential new avenue for the treatment of tubular disorders by an IV approach, although they have yet to be independently validated by other groups. Intriguingly, in non-human primates, unlike in mice, effective PT transduction can be achieved via RP injection using not only AAV-KP1 but also AAV9^12^, further complicating our understanding of AAV vector-mediated renal gene delivery. These striking variabilities in outcomes across research groups and experimental settings have been confounding and puzzling, and the difficulty in reconciling them reflects our incomplete understanding of the underlying mechanisms. Therefore, elucidating these mechanisms is crucial for overcoming current barriers and advancing the development of effective renal gene therapies.

While reviewing the work by us and others on AAV-mediated gene delivery to the kidney, we noticed unambiguous discrepancies in the relative ease of mediating PT transduction and its microscopic patterns between AAV vectors expressing monomeric fluorescent protein reporters such as green fluorescent protein (GFP) and mCherry (∼27 kDa)^21, 23, 26, 27, 28, 29^ and those expressing tandem dimer Tomato (tdTomato, ∼54 kDa)^12, 15, 16^. We therefore reasoned that a direct head-to-head comparison of AAV vectors encoding monomeric versus dimeric fluorescent proteins would provide critical insights into the mechanistic basis of these confounding discrepancies in PT transduction efficiency with AAV vectors. With these backdrops, in this study, we conducted a series of experiments in mice to understand potential mechanistic differences causing these contradicting observations using the following three fluorescent reporters, GFP (monomeric), mCherry (monomeric) and tdTomato (dimeric). As a result, we confirmed that AAV vectors expressing GFP or mCherry can mediate pseudotransduction^30^ in PTs, a phenomenon mediated by cellular uptake of proteins devoid of vector-mediated gene delivery, at exceptionally high levels that were previously overlooked and underappreciated. In addition, we elucidated its underlying mechanism of pseudotransduction in PTs, in which GFP or mCherry expressed in the liver (and potentially in other organs) by vector-mediated gene delivery is released into the circulation, pass through the glomerular filtration barrier (GFB), and are then taken up by PTs through megalin-mediated endocytosis, resulting in deterministic pseudotransduction in PTs as opposed to stochastic transduction mediated by viral vectors^31, 32^. This discovery not only fundamentally challenges the currently prevailing understanding of AAV vector-mediated PT transduction but also highlights the persistent difficulty in achieving efficient PT targeting through systemic vector delivery, even with engineered AAV capsids. Furthermore, these findings provide a framework for accurately assessing and interpreting renal transduction data and underscore the importance of considering the properties of reporter and therapeutic proteins when evaluating AAV vector-mediated gene delivery to the kidney.

## Results

### Motivation and rationale for the study

Previous studies investigating renal gene delivery with AAV vectors have reported conflicting observations in renal tubular epithelial cell transduction following administration of AAV9 vectors in mice. Following systemic or local injection of AAV9 vectors, some studies, which we term the “Uniform” group, reported effective transduction with strong tropism in renal tubular epithelial cells^21, 22, 23, 28^, whereas others, including our own, which we term the “Mosaic” group, have observed only limited transduction in renal tubular epithelial cells including PTs^12, 13, 16, 17, 18, 19^. Careful reviews of histological images of the kidneys in both the Uniform and Mosaic groups available in publications and in our own laboratory revealed features that are consistent within each group but clearly distinct between them. The Uniform group studies employed either GFP or mCherry, and observed PT signals that are predominantly localized to the luminal side of renal tubular epithelial cells, which represents an unusual subcellular distribution of cytoplasmic fluorescent reporters. In addition, the signals are contiguous along the entire tubular structure unit with no intervening non-transduced cells, and frequently exhibit a punctate pattern (**Fig. 1a**). In contrast, the Mosaic group studies used either the dimeric fluorescent protein, tdTomato, or a Cre transgene combined with the loxP-STOP-loxP-fluorescent reporter mouse strains, Ai14 mice^33^ or mT/mG mice^34^. These studies observed the expected distribution of diffuse cytoplasmic fluorescent signals in renal tubule epithelial cells with tdTomato-expressing vectors or in AAV-Cre-injected Ai14 mice, and plasma membrane-restricted signals in AAV-Cre-injected mT/mG mice. In addition, the signals are often non-contiguous with non-transduced cells interspersed within the same tubular structure unit, exhibiting a mosaic pattern (**Fig. 1b**). Given that monomeric fluorescent proteins can be filtered through glomeruli and subsequently reabsorbed by PTs, where they accumulate on the luminal side of tubular cells^35, 36^, we hypothesized that GFP and mCherry signals previously interpreted as AAV vector-mediated PT transduction might represent primarily pseudotransduction^30^ devoid of vector genome delivery to PTs, rather than bona-fide vector-mediated gene delivery. Supporting this notion, we could readily detect circulating GFP in the sera of C57BL/6J mice collected 2 weeks after IV injection of 4 x 10^13^ vg/kg body weight of AAV9-CAG-nlsGFP (n=3 in Group 1 in **Table 1**) under a fluorescence microscope (**Fig. 1c**). The kidneys harvested 2 weeks post-injection exhibited GFP signals consistent with the pattern observed in the Uniform group (**Fig. 1d** and **Supplementary Fig. 1**), even though GFP has a nuclear localization signal (nls or NLS). This strongly supported our hypothesis.

**Fig. 1.**
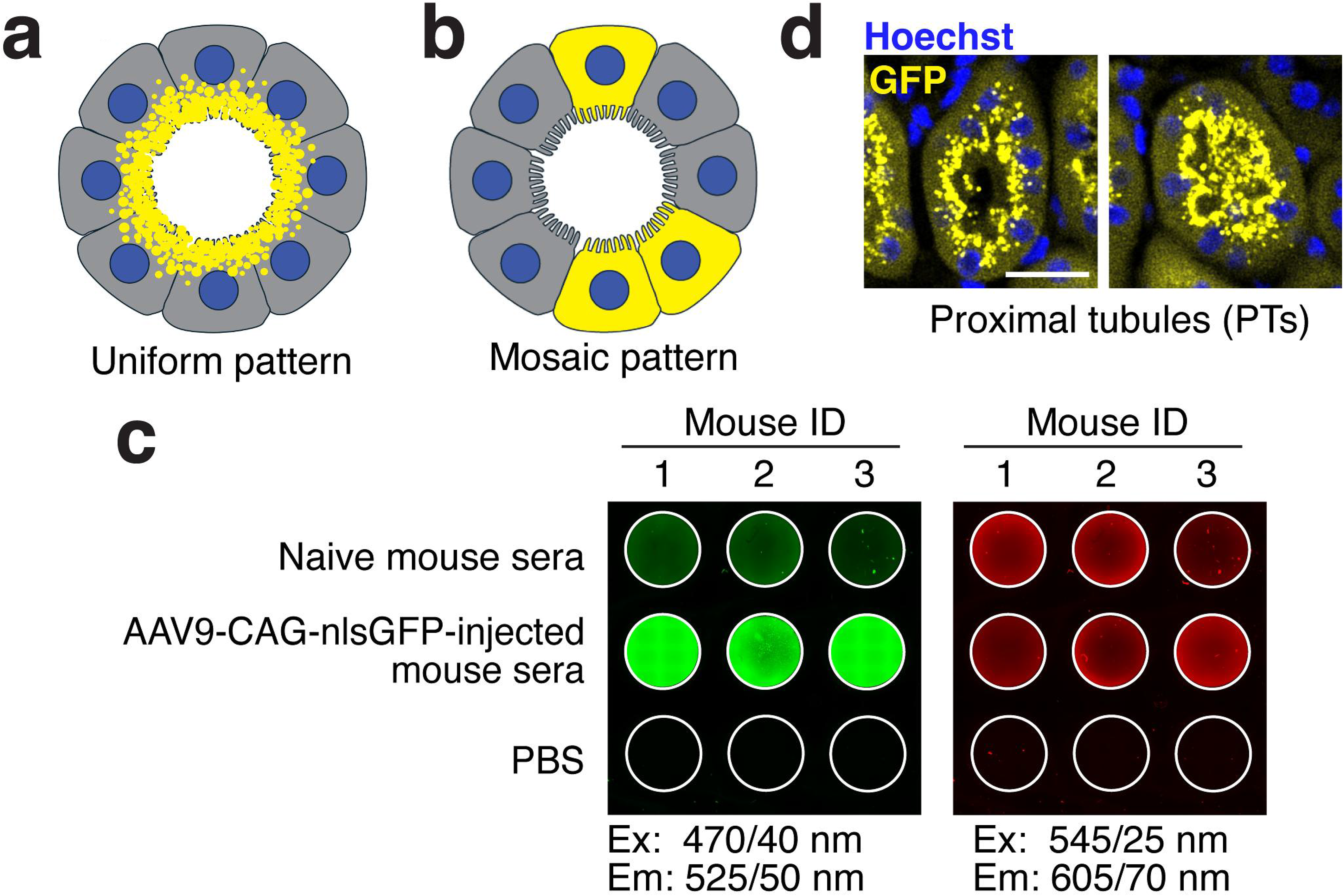
Schematic illustrating two distinct patterns of reported PT transduction and evidence of circulating GFP. Schematic transverse sections of a proximal tubule (PT) illustrating the microscopic features of fluorescent protein signal distribution observed in two distinct PT transduction patterns: the “Uniform” group (**a**) and the “Mosaic” group (**b**). In the “Uniform” group, fluorescent protein signals (yellow, *e.g.*, mCherry) form a continuous ring composed of numerous puncta located on the brush border, which is LTL-positive, and the subluminal cytoplasm in all epithelial cells surrounding the PT lumen. In contrast, the “Mosaic” group shows patchy, discontinuous fluorescent protein signals (yellow, *e.g.*, tdTomato) with diffuse intracellular localization, distinct cell boundaries, and intervening signal-negative cells. (**c**) Representative fluorescence images of serum samples observed in green and red channels. Middle row: serum from C57BL/6J male mice (n=3 in Group 1) collected 2 weeks after intravenous injection of AAV9-CAG-nlsGFP (injected at 4 x 10^13^ vg/kg body weight at 8 weeks of age). Top row: serum from age-matched uninjected C57BL/6J male mice (n=3). Bottom row: PBS controls (n=3). Images were captured using a Keyence BZ-X710 microscope (Green channel: excitation (Ex), 470/40 nm; emission (Em), 525/50 nm. Red channel: Ex, 545/25 nm; Em, 605/70 nm). Image intensities were adjusted to optimize visualization of positive signals; intensities are relative within each channel and do not reflect absolute quantification. GFP signals were not visible to the naked eyes under ambient light but were readily detectable by fluorescence microscopy. **(d)** Representative proximal tubules (PTs) from kidneys of these three mice harvested 2 weeks post-injection. Shown are from the Group 1 Mouse 2 kidney presented in **Supplementary Fig. 1**. GFP signals shown in yellow exhibited a pattern consistent with that observed in the Uniform group. PTs were identified based on green autofluorescence. Scale bar, 10 μm.

**Table 1.**
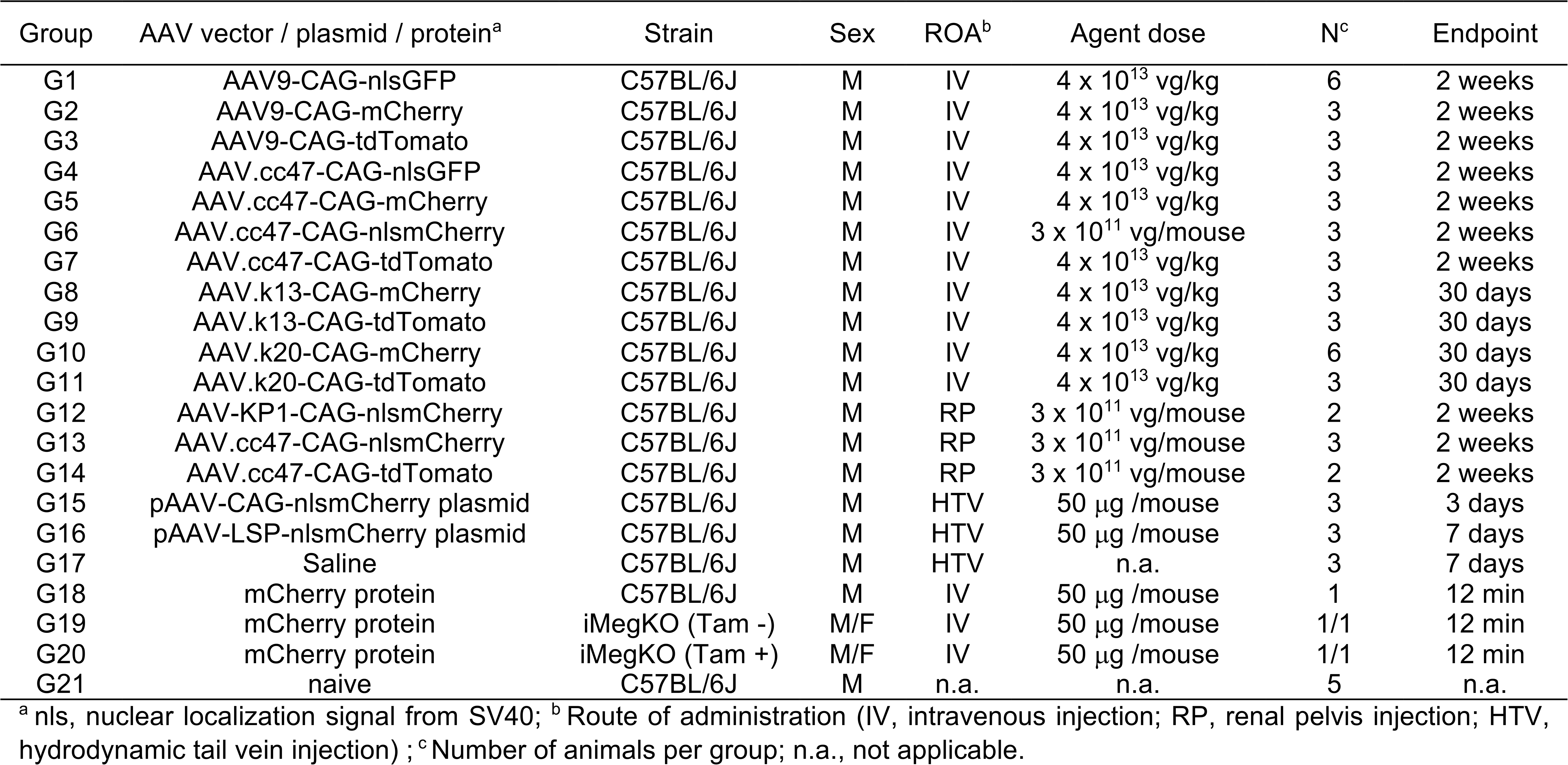
Summary of the experiments addressing AAV vector-mediated true- and pseudo-transduction in proximal tubules in mice.

### Experimental design

To rigorously test the hypothesis that the PT signals observed with GFP and mCherry fluorescent protein reporters largely reflect pseudotransduction rather than bona fide AAV-mediated gene delivery and fully understand true PT transduction profiles, we designed a set of comparative experiments using diverse fluorescent protein reporters (nlsGFP, mCherry, nlsmCherry, and tdTomato), AAV capsids (AAV9, AAV-KP1, AAV.cc47, AAV.k13, and AAV.k20), and delivery routes (IV and RP), as well as control constructs including a purified mCherry protein and plasmid vectors expressing nlsmCherry. To drive the expression of reporters from vectors, we used the CAG promoter, a strong ubiquitous promoter active in a wide range of cell types. In addition, we used a liver-specific promoter (LSP) in a plasmid construct to restrict expression of fluorescent protein reporters only in the liver. We administered these agents to 8-week-old C57BL/6J male mice and assessed the corresponding fluorescent protein signals in the kidney, liver and heart as summarized in **Table 1**. We also collected serum samples at the study endpoint. In Group 10 mice, we also collected serum samples before injection. We included fluorescent protein reporters with an NLS to conveniently distinguish true transduction events from protein uptake leading to pseudotransduction. Since proteins bound to cell surface and subsequently internalized are not trafficked to the nucleus even if they contain an NLS^37^, the nuclear enrichment of fluorescent protein reporters with an NLS provides evidence for bona fide vector-mediated transduction. GFP and mCherry are widely used monomeric fluorescent reporters, while tdTomato is a dimeric reporter twice their size. AAV9 and AAV-KP1 are contrastive capsids in renal gene delivery in mice, with the former capsid not being able to transduce PTs irrespective of route of administration when tdTomato is used, while the latter demonstrating effective PT transduction when delivered locally^12^. AAV.cc47, AAV.k13 and AAV.k20 are capsids that have been reported to effectively transduce PTs when AAV vectors expressing mCherry are delivered by IV injection^26, 27^.

### Conventional fluorescent protein reporters circulate in vector-treated mice: a prevalent but underreported phenomenon

A previous study has shown that engineered fluorescent proteins containing a secretory signal sequence, when expressed in the liver via AAV vector-mediated gene delivery, can be efficiently secreted into the circulation and readily detected for imaging applications such as visualizing the microvasculature in mice^38^. However, it has not been widely appreciated that conventional, non-engineered fluorescent proteins lacking a secretory signal can also be released into the bloodstream at appreciable levels following vector delivery. This may raise the question of whether our observation of GFP in the sera of AAV9-CAG-nlsGFP-injected mice (**Fig. 1c**) represents a general phenomenon or an exceptional case. To address this question, we investigated the presence of GFP, mCherry, and tdTomato in the circulation in vector-treated mice. Serum samples were collected at study endpoints and analyzed by western blot using the corresponding antibodies. The presence of fluorescent proteins were clearly demonstrated in all serum samples from mice injected with AAV9-CAG-nlsGFP (Group 1), AAV.cc47-CAG-nlsGFP (Group 4), AAV9-CAG-mCherry (Group 2), AAV.cc47-CAG-mCherry (Group 5), AAV.k13-CAG-mCherry (Group 8), and AAV.k20-CAG-mCherry (Group 10) (**Fig. 2a** and **Supplementary Fig. 2a, b, c, d**). Western blots of sera from mice injected with AAV9-CAG-tdTomato (Group 3), AAV.cc47-CAG-tdTomato (Group 7), AAV.k13-CAG-tdTomato (Group 9), AAV.k20-CAG-tdTomato (Group 11) revealed no bands, indicating the absence of circulating tdTomato although the presence of tdTomato at levels below the sensitivity of this assay could not be ruled out (**Supplementary Fig. 2e, f**).

**Fig. 2.**
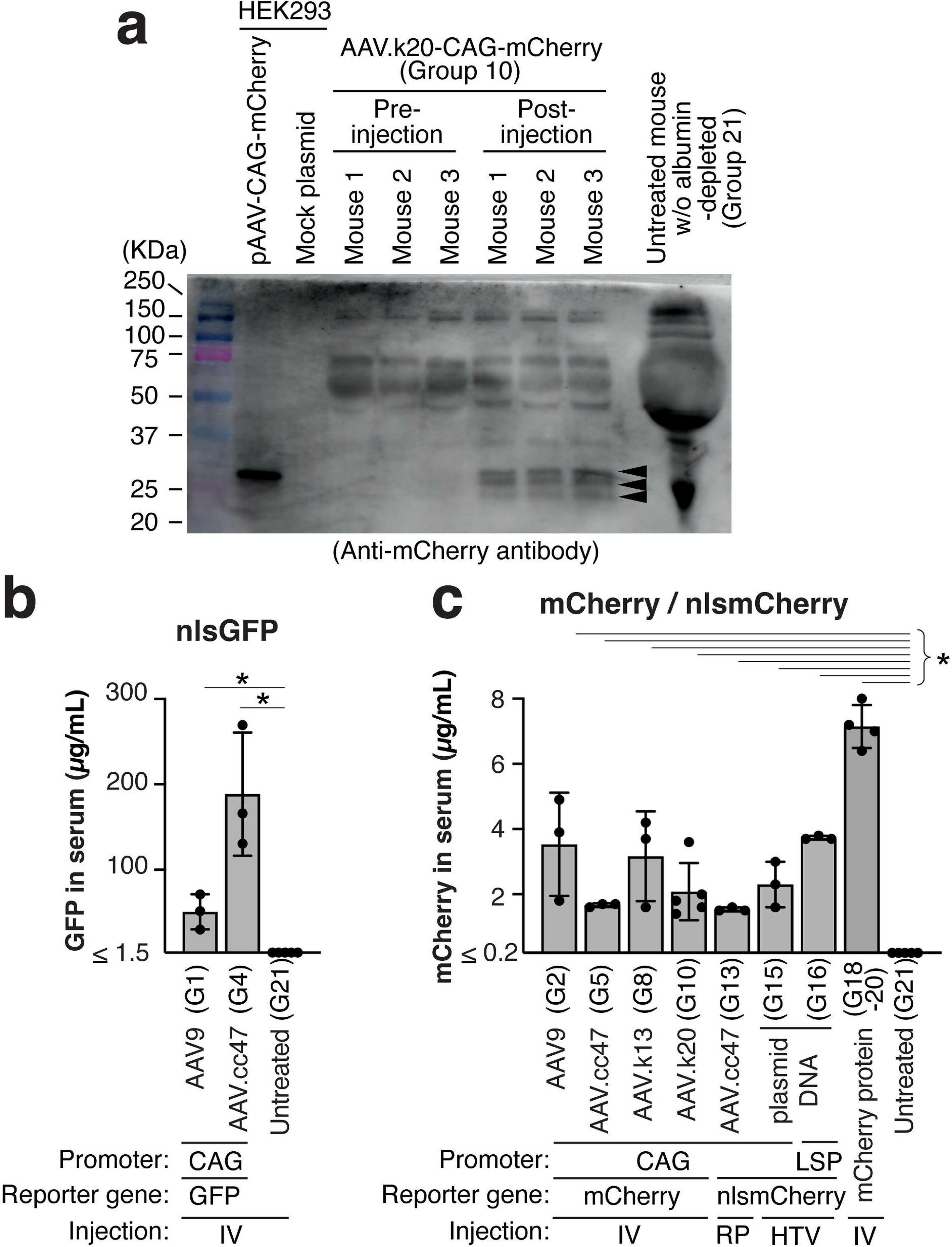
Demonstration of circulating GFP and mCherry fluorescent proteins in the blood in mice receiving AAV-GFP or AAV-mCherry vectors by western blot and serum fluorometric analyses. Serum samples from mice injected with AAV or plasmid vectors expressing fluorescent proteins were subjected to western blot analysis (Panel a) or quantitative fluorometric assay to demonstrate the presence of vector-derived fluorescent proteins in the circulation (Panels b and c). **(a)** Albumin-depleted serum samples collected before and 2 weeks after intravenous injection of AAV.k20-CAG-mCherry at 4 × 10^13^ vg/kg (Mouse 1, 2, and 3 in Group 10) were separated in an 8-16% SDS polyacrylamide gel along with naive mouse serum sample that did not undergo the albumin depletion procedure and subjected to western blot analysis with anti-mCherry antibody. Lysates of HEK293 cells transfected with pAAV-CAG-mCherry plasmid DNA and non-transfected HEK293 cell lysates were used as positive and negative controls. Arrowheads indicate mCherry protein and mCherry-derived truncated forms. **(b)** Concentrations of circulating fluorescent proteins (µg/mL) in sera from mice administered with the agents indicated in the panel were determined by the quantitative fluorometric assay we have devised (**Supplementary** Fig. 3). In this assay, serum samples collected at study endpoints (see **Table 1**) were diluted 7.5-fold in PBS, and fluorescence intensity was measured using a PerkinElmer Victor Nivo^TM^ Multimode Plate Reader with the excitation/emission filters: 480/530 nm for GFP and 480/530 nm for mCherry. Sera from 5 naive untreated C57BL/6J mice served as negative controls. Data are presented as mean ± SD with individual data values shown as dots. Statistical comparisons between each agent-treated group and the naive control group were performed using two-tailed Mann-Whitney U test with Benjamini-Hochberg FDR adjustment. Asterisks indicate statistical significance with α=0.05. q values are all 0.0179.

To further confirm the presence of fluorescent proteins in the circulation in fluorescent protein-expressing AAV vector-injected mice, we developed a fluorometric assay that can quantify GFP, mCherry and tdTomato proteins present in mouse sera. Based on four-parameter logistic regression analysis, the dynamic ranges of the developed assays were determined to be: 1.5 to 750 µg/mL in mouse serum for GFP, 0.2 to 750 µg/mL in mouse serum for mCherry, and 2.9 to 750 µg/mL in mouse serum for tdTomato (**Supplementary Fig. 3,** and **Supplementary Data 1**). Using this quantitative fluorometric assay, we investigated the presence of AAV vector-derived fluorescent proteins in sera collected from GFP-, mCherry-, nlsmCherry- and tdTomato-expressing AAV vector-injected mice (Groups 1, 2, 3, 4, 5, 6, 7, 8, 9, 10, 11, and 13). Untreated naive mouse sera (Group 21, n=5) served as negative controls. For the assay for mCherry, we also included sera collected 12 min post-injection of purified mCherry protein (Groups 19 and 20) and sera collected from pAAV-CAG-nlsmCherry plasmid-injected mice (Groups 15) and pAAV-LSP-nlsmCherry plasmid-injected mice (Group 16), which were obtained in the experiments described below. These assays confirmed the presence of GFP and mCherry in sera from nlsGFP-, mCherry-, or nlsmCherry-expressing vector-injected mice at levels of 30 to 269 µg/mL for nlsGFP and 1.4 to 4.9 µg/mL for mCherry and nlsmCherry (q=0.0179, two-tailed Mann-Whitney U test with Benjamini-Hochberg FDR adjustment) (**Fig. 2b, c**), while tdTomato was detected in none of the sera from tdTomato vector-injected mice with our assay with the lower limit of detection of 2.9 µg/mL. Taken together, these data provide evidence that GFP and mCherry proteins, even in the absence of a secretory signal, can be released into the bloodstream from transduced organs and circulate systemically in mice injected with AAV vectors expressing a fluorescent protein reporter.

### AAV vectors expressing GFP and mCherry mediate misleading pseudotransduction in PTs

Kidneys, livers and hearts were harvested from mice injected intravenously with AAV vectors expressing nlsGFP (Groups 1 and 4), mCherry, or nlsmCherry (Groups 2, 5, 6, 8, and 10) at study endpoints (2 weeks or 30 days post-injection, please refer to **Table 1**). Kidney tissues were subjected to immunofluorescent microscopic analyses using both epifluorescence and confocal laser scanning microscopes to observe nlsGFP, mCherry, or nlsmCherry signals together with PT-specific markers, Lotus tetragonolobus lectin (LTL), and megalin coded by the *LRP2* gene (**Fig. 3** and **Supplementary Fig. 4**). LTL specifically binds to fucose-containing glycoconjugates present on the brush border of PT epithelial cells. Megalin is an endocytic receptor that localizes to the luminal surface of PT epithelial cells, where it mediates the reabsorption of various protein^39^. Please note that we used fluorescein-conjugated LTL and it was used only for mCherry and nlsmCherry samples because of significant spectral overlap between GFP and fluorescein. Regardless of whether the capsid is AAV9, AAV.cc47, AAV.k13, or AAV.k20, and regardless of the presence or absence of an NLS in the fluorescent protein reporters, we observed the same pattern of reporter signals strongly associated with megalin signals throughout the renal cortex and the outer medulla (**Fig. 3** and **Supplementary Fig. 4**). Both mCherry and nlsmCherry signals were significantly enriched at the inner boundary of LTL signals, where the fluorescent reporter and megalin signals overlapped and exhibited a punctate pattern. nlsGFP signals were similarly enriched on the luminal surface and in the subluminal cytoplasmic region. Importantly, no nuclear signals in PTs were observed with either nlsGFP or nlsmCherry. These observations were inconsistent with the pattern exhibited by nlsGFP or nlsmCherry synthesized in cells as a consequence of true AAV vector transduction or the “Mosaic” pattern (**Fig. 1b**) that we anticipated based on our previous observations of true AAV vector-mediated PT transduction^12, 15^.

**Fig. 3.**
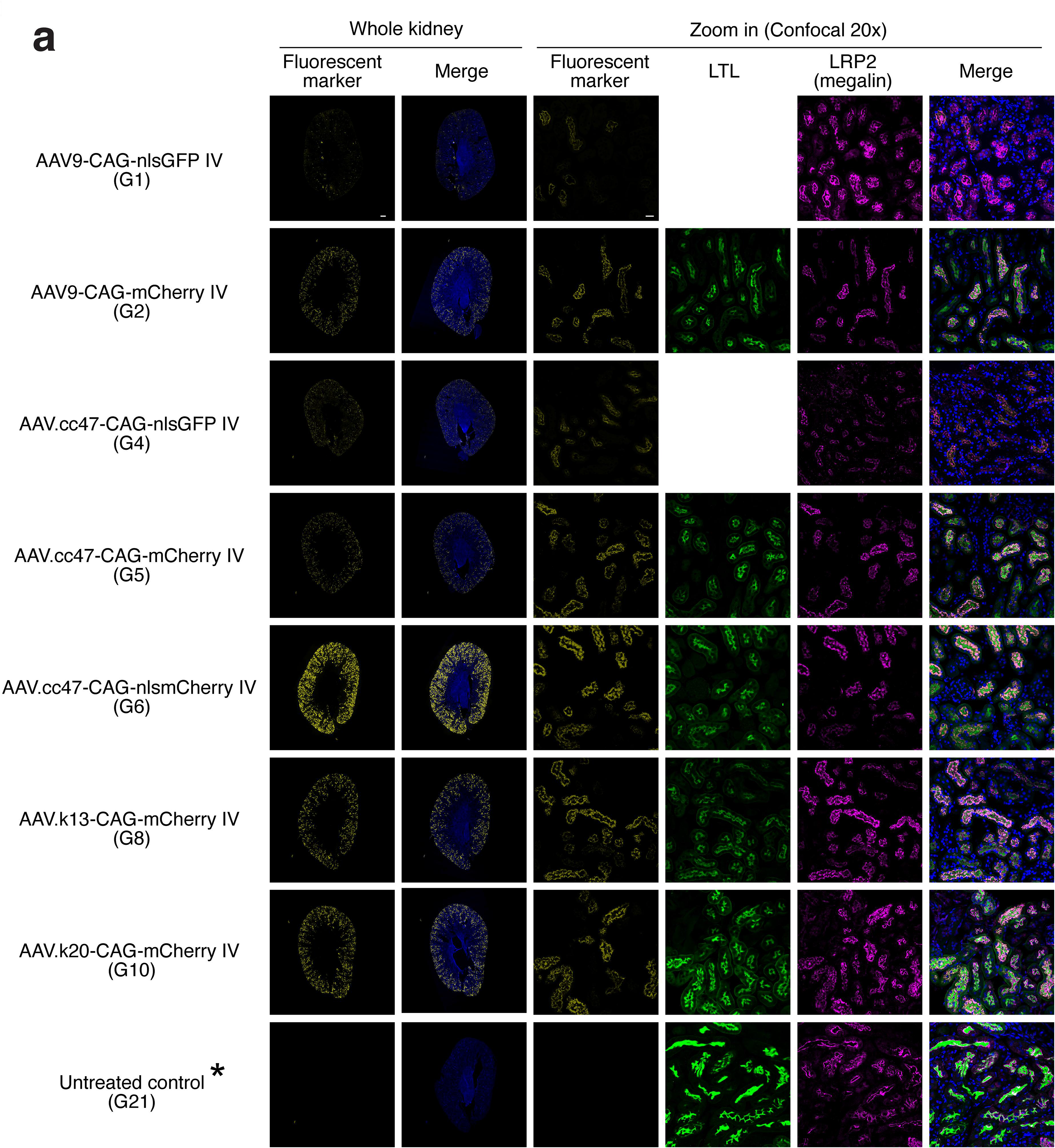
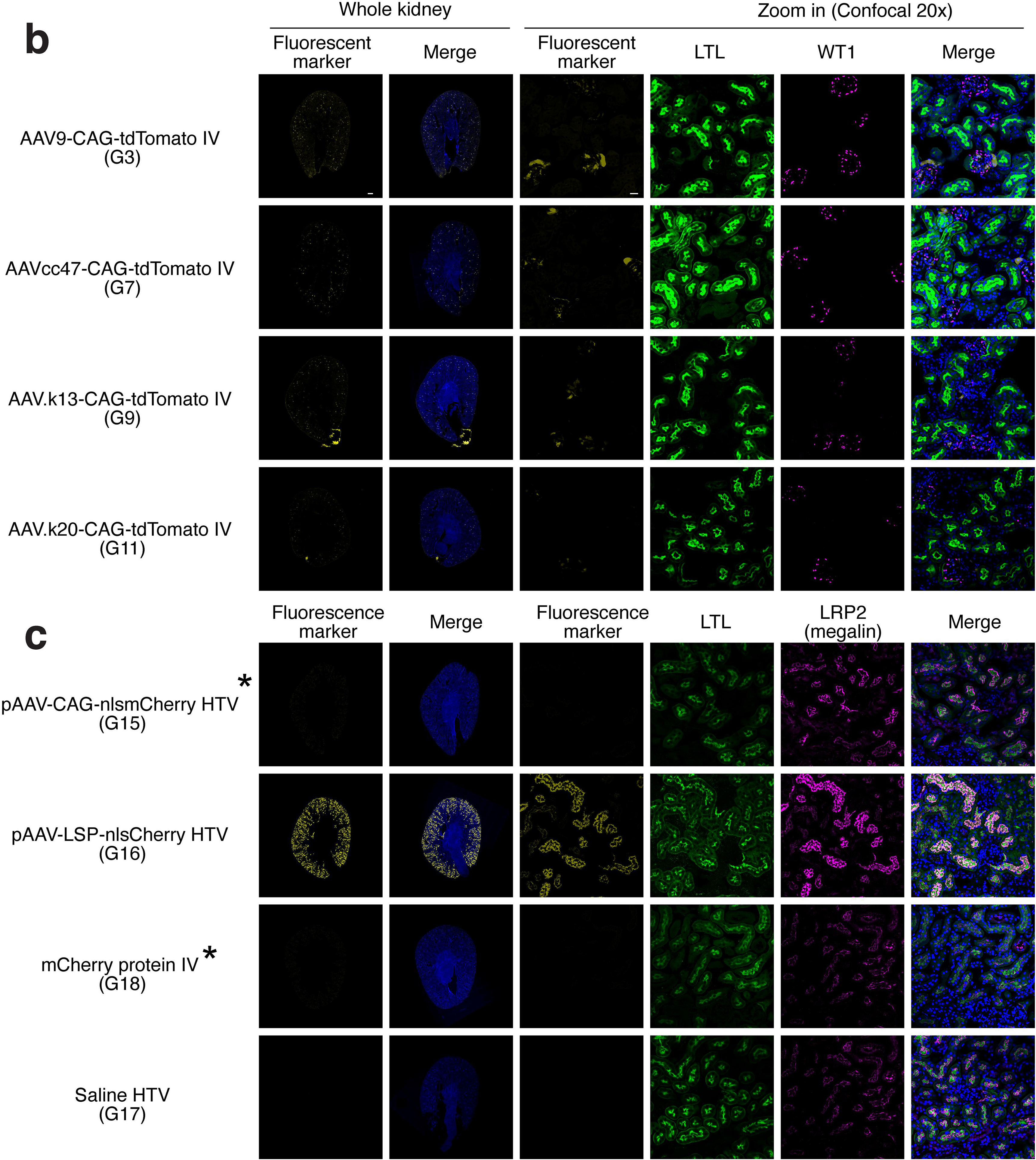
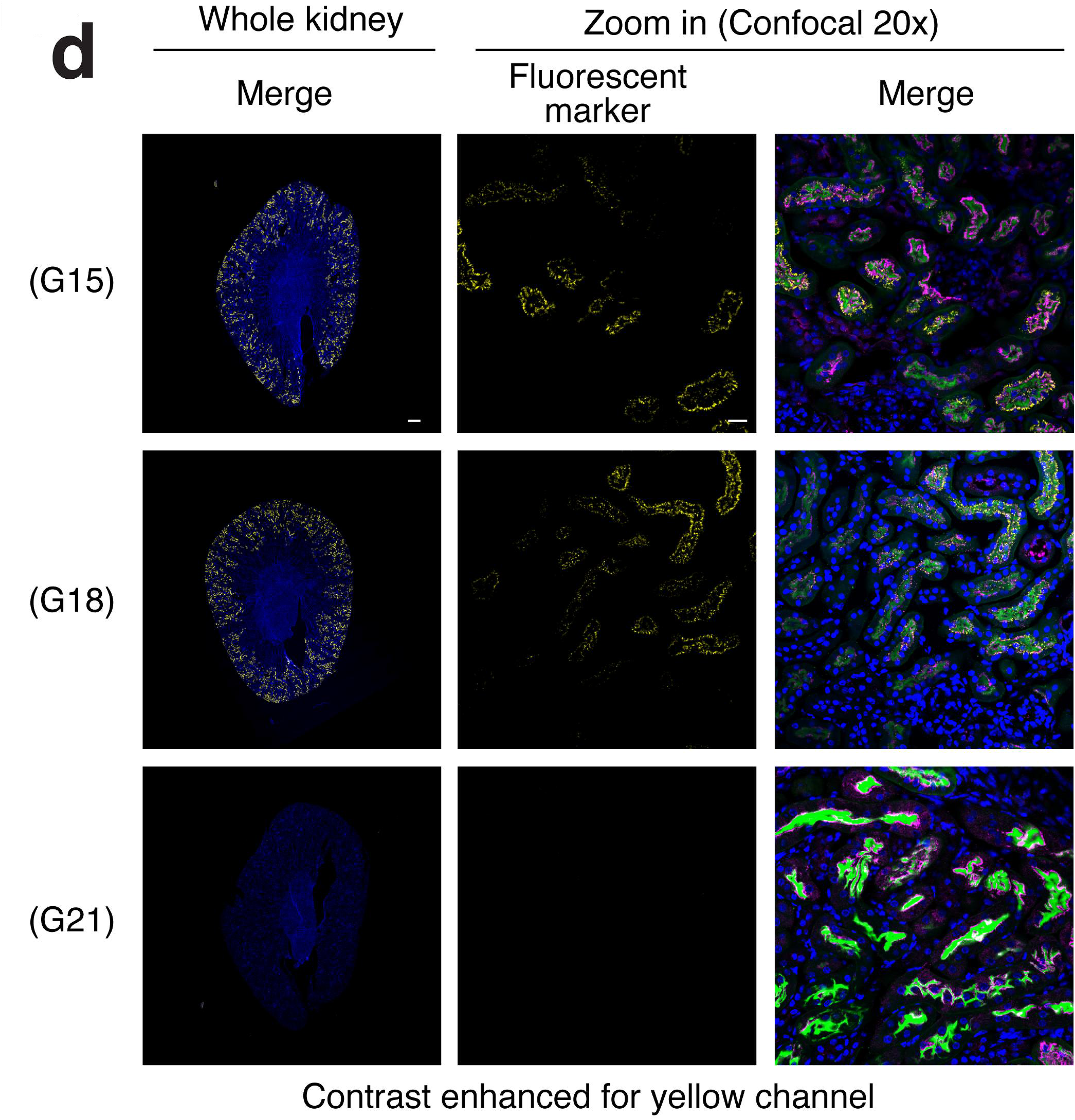
Immunofluorescence microscopy images of fluorescent protein signals in the kidneys of mice receiving viral and plasmid vectors expressing fluorescent reporters and purified fluorescent protein agents. Kidneys harvested at each study endpoint were subjected to immunofluorescence microscopy using a Keyence BZ-X710 epifluorescence microscope and an LSM 900 confocal microscope. Images show fluorescent reporter signals (yellow) and marker signals obtained with Lotus tetragonolobus lectin (LTL, green), anti-LRP2 (megalin) or anti-WT1 antibody (magenta), and Hoechst 33342-stained nuclei (blue). **(a)** Representative whole kidney epifluorescence images (left panels) and high-magnification confocal images (right panels) of kidney sections from mice injected intravenously with AAV vectors expressing either nlsGFP, mCherry, or nlsmCherry. Untreated kidney (Group 21) serves as negative controls showing no appreciable bleed-through of megalin signals in the fluorescent reporter channel. Vector and group number information (see **Table 1**) is indicated for each set of images. Contrast enhanced images of the groups indicated with an asterisk are shown in Panel c. The absence of nuclear localization despite NLS in nlsGFP and nlsmCherry fluorescent markers, and the enrichment of fluorescent signals at the luminal surface and in the subluminal space with punctate patterns showing substantial overlap with megalin demonstrate that the observed signals represent pseudotransduction. **(b)** Representative whole kidney images and high-magnification confocal images of kidney sections from mice injected intravenously with AAV vectors expressing tdTomato. Signals are primarily from glomeruli (mesangial cells) and occasionally from the thick ascending limb (TAL) of the loop of Henle (TAL) and interstitial cells. Signals from PTs could barely be observed. The absence of PT signals has been confirmed with contrast-enhanced images (**Supplementary Fig. 5**). **(c)** Representative whole kidney images and high-magnification confocal images of kidney sections from mice receiving hydrodynamic tail vein (HTV) injection of plasmid DNAs expressing nlsmCherry or saline (a negative control) or intravenous injection of purified mCherry protein. Any signals observed in the kidneys in these mice represent signals originating from circulating mCherry protein. **(d)** Additional whole kidney and confocal images in Groups 15, 18 and 21 demonstrating the presence (in G15 and G18) and absence (in G21) of mCherry signals in PTs. Yellow channel contrast was enhanced by linearly adjusting signal intensities. “Merge” indicates images showing all the colors shown in the panels together with Hoechst-stained nuclei. Scale bars are indicated in the images (300 µm for the whole kidney images and 20 µm for confocal images, shown only in the top row). High-resolution raw and processed images have been deposited in Zenodo. Image display settings are summarized in **Supplementary Table 3**.

Liver sections were also examined by epifluorescence and confocal laser scanning microscopy. All the liver sections in Groups 1, 2, 4, 5, 6, 8, and 10 demonstrated hepatocyte transduction at varying levels, with enrichment around the central vein (**Fig. 4** and **Supplementary Fig. 5**). The expected nuclear-enriched signals in hepatocytes were observed in mice injected with the nlsGFP- or nlsmCherry-expressing vectors (Groups 1, 4 and 6) (**Fig. 4** and **Supplementary Fig. 5**), while liver sections from mCherry-expressing vectors (Groups 2, 5, 8, and 10) demonstrated a characteristic pattern of signal distribution in hepatocytes comprising two distinct compartments: the diffuse cytoplasmic signals and intense punctate signals (**Fig. 4** and **Supplementary Fig. 5**). The punctate pattern of mCherry and other monomeric red fluorescent proteins has been documented in previous studies and reflects their resistance to acidic quenching and lysosomal degradation^40, 41^. Thus, the observed mixed pattern of diffuse and punctate signals is consistent with overexpression of cytosolic mCherry protein mediated by AAV vector transduction in hepatocytes. Heart sections were also examined by epifluorescence microscopy. While showing varying levels of transduction in the heart, all the AAV vector-injected groups exhibited the anticipated cardiac transduction patterns with cytosolic and nuclear-enriched fluorescent reporters (**Fig. 4** and **Supplementary Fig. 5**).

**Fig. 4.**
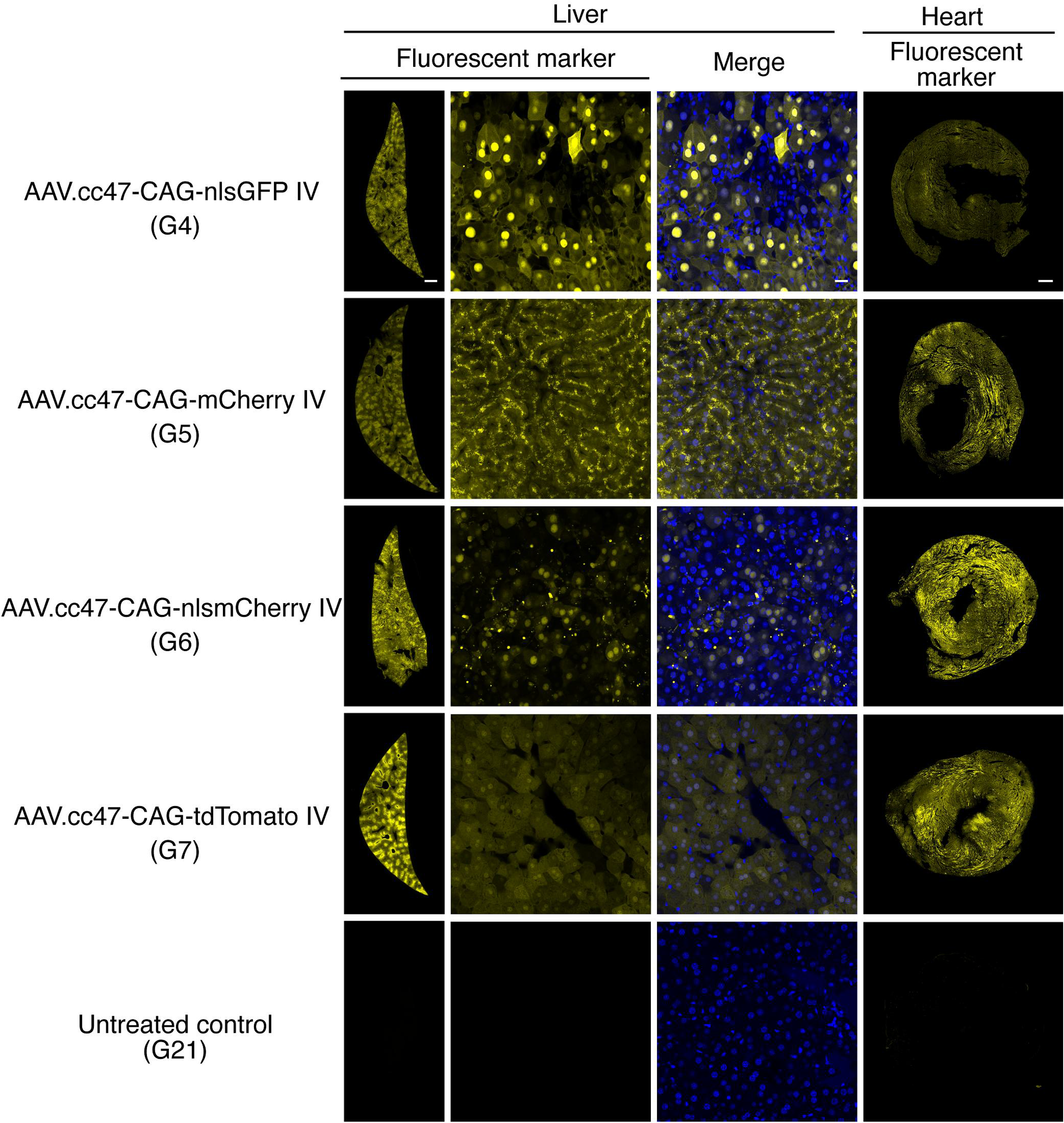
Liver and heart transduction in mice following IV administration of AAV.cc47 vectors expressing different fluorescent reporters. Representative epifluorescence and confocal microscopy images of liver and heart sections from mice in Groups 4, 5, 6 and 7, injected intravenously with AAV.cc47 vectors expressing different fluorescent reporters and in uninjected control group (Group 21). The images in the leftmost and rightmost columns represent the whole liver and heart sections, respectively, showing only fluorescent marker signals. The middle-left liver confocal images show only fluorescent marker signals, while the middle-right images include Hoechst-stained nuclear signals. AAV.cc47-CAG-nlsGFP (Group 4) and AAV.cc47-CAG-nlsmCherry (Group 6) mediated hepatocyte transduction with expected nuclear-enriched GFP and mCherry signals. AAV.cc47-CAG-tdTomato (Group 7) generated expected diffuse signals in hepatocytes. In contrast, AAV.cc47-CAG-mCherry (Group 5) mediated hepatocyte transduction exhibiting the characteristic mixed pattern of diffuse cytoplasmic and punctate signals in hepatocytes. Untreated control (Group 21) shows absence of fluorescent signals. All AAV.cc47 vector-treated groups show cardiac transduction at varying levels. Scale bars in the top row represent 1 mm, 20 µm, and 300 µm from left to right. High-resolution raw and processed images have been deposited in Zenodo. High-resolution raw and processed images have been deposited in Zenodo. Image display settings are summarized in **Supplementary Table 3**.

With all these observations, we have concluded that the observed fluorescent reporter signals in PTs in the groups receiving intravenous injection of AAV vectors expressing GFP or mCherry primarily represent pseudotransduction.

### Circulating mCherry protein leads to pseudotransduction in PTs

To obtain direct evidence that circulating mCherry protein leads to pseudotransduction in PTs, we injected 50 µg of affinity column-purified mCherry protein intravenously into one 8-week-old C57BL/6J male mouse and harvested the kidneys 12 min post-injection (Group 18). We confirmed the presence of circulating mCherry protein following injection in this animal (**Fig. 2c**). Histological analysis of the kidney revealed mCherry signals in PTs that mirrored the “Uniform” pattern (**Fig. 1a**), although the signals were much weaker than those in AAV vector-injected mice (**Fig. 3c, d**, and **Supplementary Fig. 4a**) likely due to the transient exposure of PTs to mCherry in contrast to the sustained exposure achieved through AAV vector-mediated pseudotransduction in PTs. To further strengthen the evidence that pseudotransduction in PTs is mediated by circulating mCherry protein, we injected 50 µg of the pAAV-CAG-nlsmCherry (Group 15) or pAAV-LSP-nlsmCherry plasmid (Group 16) into 8-week-old C57BL/6J male mice using the hydrodynamic tail vein (HTV) injection technique^42^, and harvested the kidney and liver tissues 3 and 7 days post-injection for Group 15 and Group 16, respectively. Although the HTV technique can transfect hepatocytes in the liver with injected DNA effectively (**Supplementary Fig. 5**), DNA delivery to the kidney is minimal^43, 44^. In both Group 15 and Group 16 mice, circulating nlsmCherry protein in sera was detected at levels comparable to the AAV-mCherry/nlsmCherry vector-injected mice at the study end points (3 and 7 days post injection in Group 15 and Group 16, respectively) (**Fig. 2c**). Histological analysis of the kidneys revealed much stronger mCherry signals in PTs exhibiting the “Uniform” pattern (**Fig. 1a**) than the protein-injected group (Group 18) (**Fig. 3c, d**, and **Supplementary Fig.4a**) likely due to more sustained circulation of mCherry in the blood in the plasmid-injected groups. Since de novo mCherry protein synthesis in PTs can be confidently excluded in the protein and plasmid-injected groups (Groups 15, 16, and 18), we concluded that the observed pseudotransduction in PTs is mediated by circulating mCherry protein. Although our study did not conclusively determine which AAV vector-transduced organs release mCherry protein into the blood, leading pseudotransduction in PTs, the data from the pAAV-LSP-nlsmCherry plasmid-injected group (Group 16) strongly indicate that the liver is one of the sources. This is because HTV-mediated hepatocyte damage is transient, with serum ALT levels returning to normal range within several days post-injection^42, 45^; therefore, nlsmCherry leakage from damaged hepatocytes in Group 16 mice should be minimal and not contribute significantly to the presence of circulating nlsmCherry protein. Thus, hepatocytes over-expressing mCherry protein, even without an NLS, can release them into the blood circulation by a not-yet-identified mechanism, leading to misleading pseudotransduction in PTs in AAV vector-injected animals. It is reasonable to infer that the same mechanism underlies pseudotransduction in PTs observed in AAV-GFP vector-injected animals.

### True PT transduction mediated by AAV vectors expressing mCherry

Although we demonstrated that GFP and mCherry fluorescent reporters could mediate misleading pseudotransduction in PTs, which may have been previously misinterpreted as true AAV vector-mediated transduction, it remained necessary to establish how authentic PT transduction with these monomeric fluorescent proteins should manifest. To address this, we injected 8-week-old C57BL/6J male mice with AAV-KP1-CAG-nlsmCherry via unilateral RP injection to the left kidney (LK) at a dose of 3 x 10^11^ vg/mouse (Group 12). The rationale for this experiment was that the demonstration of the expected true transduction pattern with an AAV-nlsmCherry vector exhibiting the “Mosaic” transduction (**Fig. 1b**) with enriched nuclear signals would further strengthen our conclusion described above. The RP route was chosen because we had not been aware of AAV capsids that can effectively mediate bona fide PT transduction by IV administration into adult wild-type C57BL/6J mice, and AAV-KP1 was selected because this capsid has been shown to effectively transduce PTs via RP injection with no ambiguity caused by pseudotransduction^12^. We also included RP injection of AAV.cc47-CAG-nlsmCherry to investigate whether this capsid can transduce PTs when delivered via RP injection (Group 13). Two weeks post-injection, both AAV vector-injected LKs and uninjected right kidneys (RKs), livers, and hearts were collected and analyzed by immunofluorescence microscopy in the same manner as described above (**Fig. 5, Supplementary Fig.4a** and **Supplementary Fig. 5**). In mice injected with AAV-KP1-CAG-nlsmCherry via RP administration, we observed, in the vector-injected LKs, sporadic mCherry signals showing the characteristic true transduction pattern; that is, nuclear-enriched signals with no signal enrichment on the luminal surface or in the subluminal cytoplasm and often together with signals representing pseudotransduction in PTs (**Fig. 5a, b**). Such nuclear-enriched signals demonstrating true transduction were observed both in LTL-positive PTs and LTL-negative cells (**Fig. 5b, c**), consistent with our previous observation that RP injection of AAV-KP1 vector transduced non-PT cells, including parietal cells, collecting duct cells, and interstitial cells^12^. It should be noted that true transduction events identifiable within pseudotransduced PTs in fluorescence microscopic images (**Fig. 5b**, arrows) likely represent only a fraction of the total events due to much stronger pseudotransduction signals that may mask true transduction. Only pseudotransduction signals with no nuclear enrichment of nlsmCherry were observed in AAV-KP1 vector-uninjected right kidneys (RKs). RP injection of AAV.cc47-CAG-nlsmCherry led only to pseudotransduction in PTs in both vector-injected LKs and uninjected RKs (**Fig. 5a** and **Supplementary Fig.4a**), indicating the inability of AAV.cc47 to transduce PTs via this route. These results demonstrate that nlsmCherry protein synthesized in cells by true transduction can correctly traffic to the nuclei within PT epithelial cells, and therefore, reasonably excludes the possibility that punctate reporter signals localized to the luminal surface and in the subluminal cytoplasm of PTs primarily represent true AAV vector-mediated transduction.

**Fig. 5.**
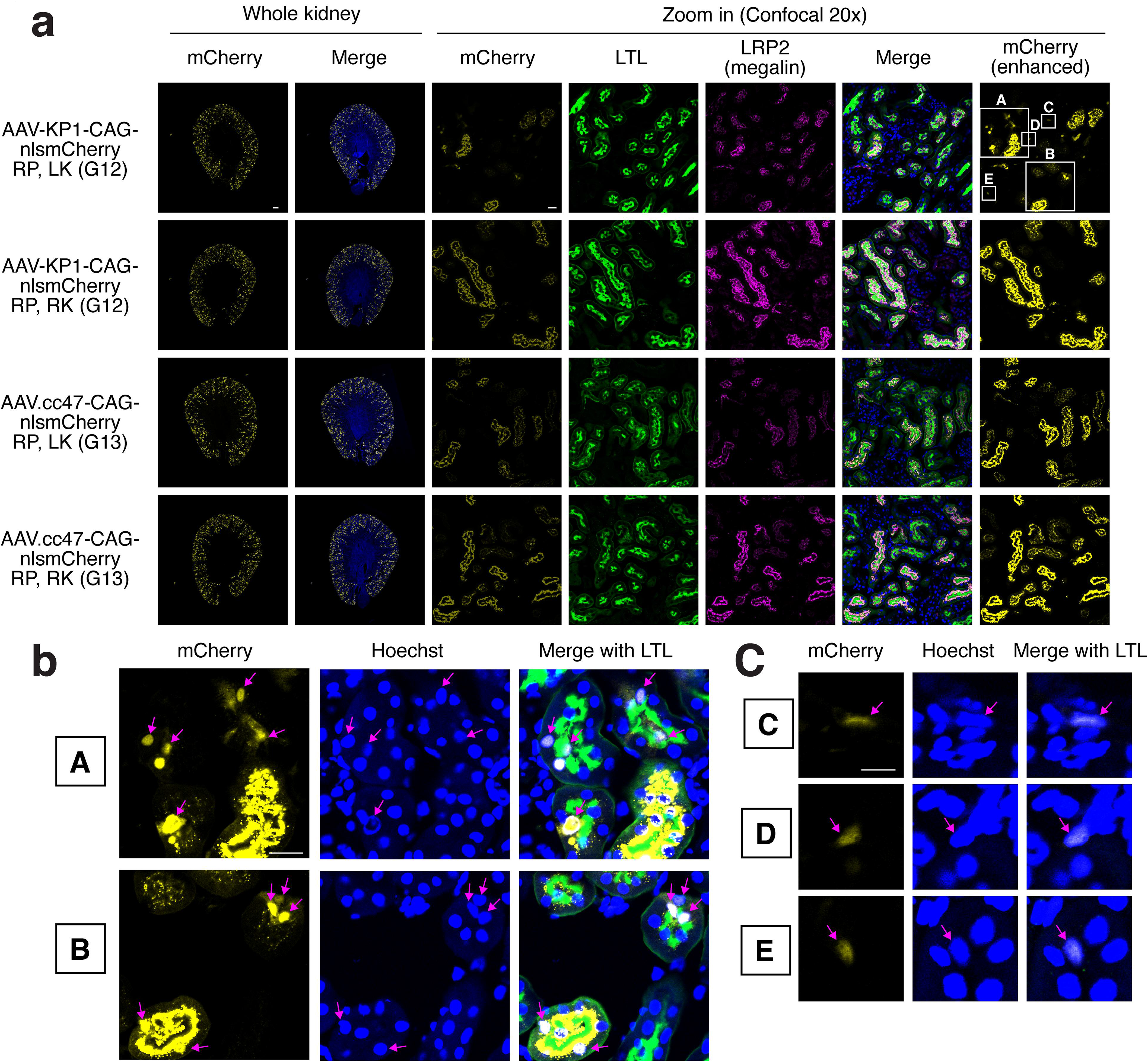
Demonstration of true transduction in PTs by RP injection of AAV-KP1-CAG-nlsmCherry in mice. **(a)** Representative whole kidney epifluorescence images (left panels) and high-magnification confocal images (right panels) of kidney sections from mice injected with AAV-KP1-CAG-nlsmCherry (Group 12) or AAV.cc47-CAG-nlsmCherry (Group 13) via unilateral renal pelvis (RP) injection into the left kidney (LK). Right kidneys (RKs) were not injected. Kidneys were harvested 2 weeks post-injection and analyzed for mCherry (yellow), LTL (green), LRP2/megalin (magenta), and Hoechst (blue) signals by immunofluorescence microscopy. Because nuclear-enriched mCherry signals representing true transduction were not as strong as the punctate luminal and subluminal mCherry signals representing pseudotransduction, contrast-enhanced mCherry images are also included to help identify true transduction. AAV-KP1-injected LK shows nuclear-enriched true transduction signals in both PTs and non-PT cells as well as pseudotransduction signals, whereas the uninjected RK displays only pseudotransduction signals. AAV.cc47-injected kidneys show only pseudotransduction signals in both LK and RK. Boxed regions (A, B, C, D, E) in the top right panel indicate areas shown at higher magnification in Panels b and c. Scale bars are indicated in the images (300 µm for the whole kidney images and 20 µm for confocal images, shown only in the top row). **(b)** High-magnification confocal images of regions A and B from Panel a, showing nuclear-enriched mCherry signals (magenta arrows) characteristic of true transduction in LTL-positive PTs within the AAV-KP1-injected LK. Please note that true transduction events can be observed in both pseudotransduced and non-pseudotransduced PTs. **(c)** High-magnification confocal images of regions C, D, and E from Panel a, demonstrating nuclear-enriched mCherry signals (magenta arrows) in LTL-negative non-PT cells in the AAV-KP1-injected LK. Scale bars are indicated in the images (300 µm for the whole kidney images and 20 µm for confocal images in Panel a, 20 µm for Panel b, and 10 µm for Panel c, shown only in the top row images in each panel). High-resolution raw and processed images have been deposited in Zenodo. Image display settings are summarized in **Supplementary Table 3**.

### Revisiting PT transduction of AAV.cc47, AAV.k13, and AAV.k20 using a dimeric fluorescent reporter, tdTomato

We have previously demonstrated that RP injection of the AAV-KP1-CAG-tdTomato vector efficiently transduces PTs in mice, whereas IV injection of the same dose of AAV9-CAG-tdTomato fails to do so. This conclusion is supported not only by the histological assessment of tdTomato signals but also by molecular evidence showing a significant difference in vector genome copy numbers between AAV9 IV and AAV-KP1 RP in the kidney (AAV9:AAV-KP1=1:26)^12^. These results indicate that when tdTomato is used as a reporter, histological observations and vector genome quantification are concordant; therefore, tdTomato signals in the kidney provide a reliable readout of true PT transduction.

On this basis, using a dimeric fluorescence reporter, tdTomato, we reassessed renal transduction profiles of AAV.c47, AAV.k13, and AAV.k20, all of which have been reported to transduce PTs effectively following IV administration when mCherry reporter was used^26, 27, 28^. Robust PT transduction by these capsids following IV delivery has also been highlighted in perspective and review articles and is emerging as a widely appreciated view on kidney transduction^46, 47, 48^. To independently verify these findings using a reporter unaffected by pseudotransduction in PTs, we injected 8-week-old C57BL/6J male mice with AAV9-CAG-tdTomato, AAV.cc47-CAG-tdTomato, AAV.k13-CAG-tdTomato and AAV.k20-CAG-tdTomato via IV at a dose of 4 x 10^13^ vg/kg (Groups 3, 7, 9 and 11, respectively).

To verify the above-described observations with RP injection of AAV.cc47-CAG-nlsmCherry, showing no true PT transduction, we also included a group receiving AAV.cc47-CAG-tdTomato via RP at a dose of 3 x 10^11^ vg (Group 14). Two weeks or 30 days post-injection, kidney tissues were collected and analyzed by immunofluorescence microscopy. This experiment revealed that all of AAV.cc47, AAV.k13, and AAV.k20 vectors transduced cells in glomeruli (mesangial cells), the thick ascending limb (TAL) of the loop of Henle, and a fewer number of cells in the interstitium, but failed to transduce PTs via IV administration (**Fig. 3, Supplementary Fig. 4b, Supplementary Fig. 6**) similar to AAV9 as we have previously reported^10, 12^. Similarly, RP injection of AAV.cc47 did not transduce PTs and resulted in the transduction of only mesangial cells, TAL cells, and interstitial cells, in both the injected and contralateral uninjected kidneys, similar to the transduction profiles observed following IV injection (**Fig. 5a, Supplementary Fig. 4b,** and **Supplementary Fig. 6**). Taken together, contrary to the previous reports, none of AAV.cc47, AAV.k13, or AAV.k20 transduced PTs in mice by IV injection, similar to AAV9. We also confirmed that RP injection of AAV.cc47 does not transduce PTs.

### Molecular evidence supporting true and pseudotransduction in PTs

Total DNA was extracted from the kidneys collected at the study endpoint in Groups 1, 2, 3, 4, 5, 7, 8, 9, 10, 11, 12 and 13, and subjected to quantitative PCR (qPCR) analysis to quantify AAV vector genome copy numbers. In these groups, immunofluorescence microscopic analyses described above demonstrated true transduction in PTs only in the kidneys receiving unilateral injection of AAV-KP1 vector (Group 12, LKs) but in none of the kidneys in the mice receiving intravenous injection of AAV9, AAV.cc47, AAV.k13 or AAV.k20 vector, vector-injected LKs or uninjected RKs in mice receiving unilateral RP injection of AAV.cc47 vector, or uninjected RKs in mice receiving unilateral RP injection of AAV-KP1 vector. In consistent with the histological observations, vector genome copy numbers in the kidneys showing both pseudotransduction and true transduction in PTs were much higher than the kidneys showing only pseudotransduction in PTs (**Fig. 6a**). Total RNA was also extracted from kidneys in Groups 1, 2, 3, 4, 5, 7, 12, and 13 and subjected to quantitative RT-PCR (RT-qPCR) analysis to quantify AAV vector genome transcript levels. The data were in full concordance with the vector genome copy number data, showing significantly elevated transcript levels only in the kidneys with true transduction in PTs (**Fig. 6c**). To enable robust statistical comparison of vector genome copy numbers and transcript levels between kidneys with and without PT transduction, Groups 12 and 13 were combined to increase statistical power. Significant differences with α=0.05 were observed for both copy numbers and transcript levels (two-tailed Mann-Whitney U test, p=0.044, **Fig. 6b, c**). These molecular data demonstrate that true transduction and pseudotransduction represent qualitatively distinct phenomena, with only the former resulting from successful AAV vector genome delivery and transgene expression in PT cells.

**Fig. 6.**
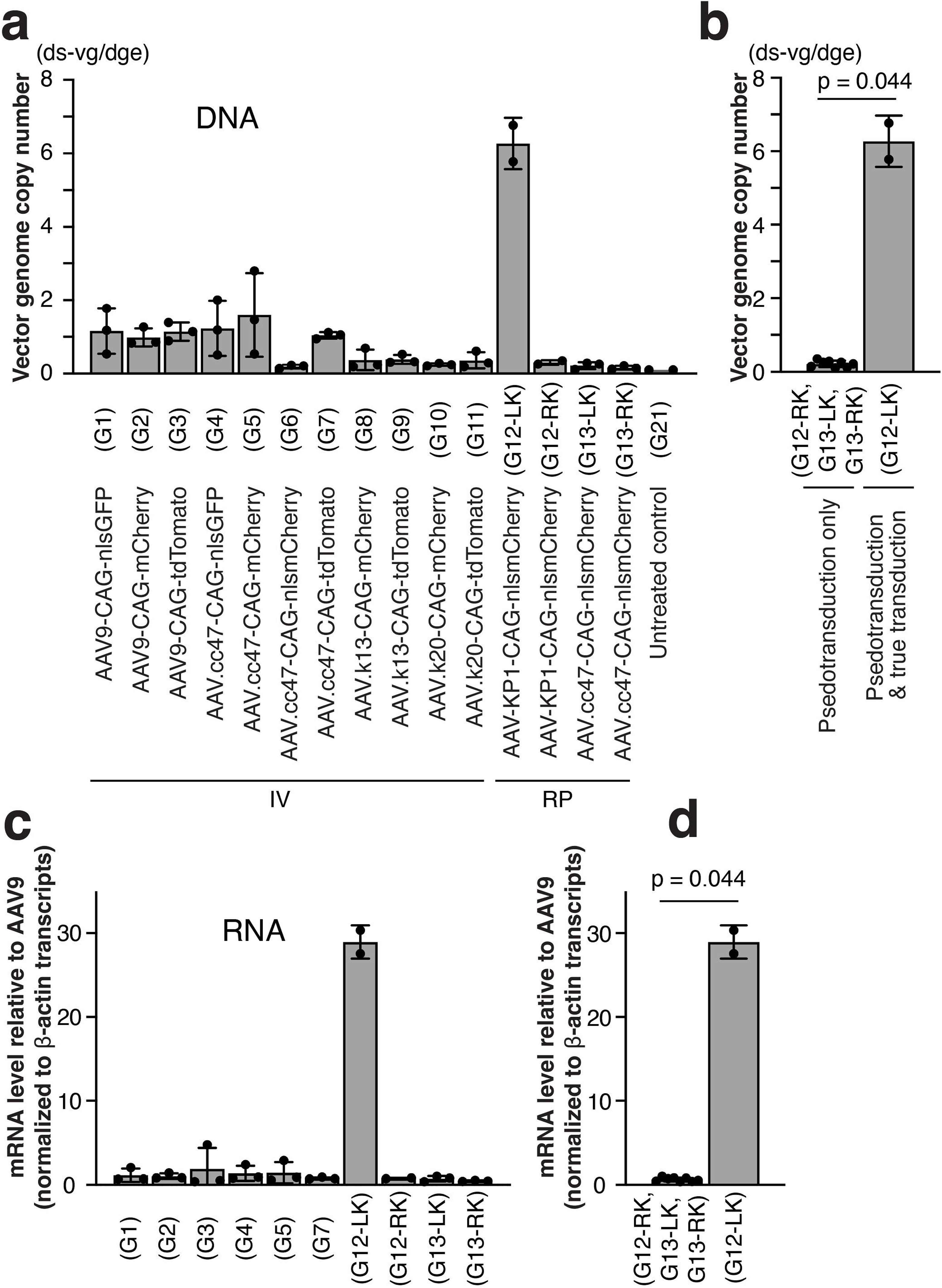
Quantification of vector genomes and transcripts differentiates pseudotransduction and true transduction in PTs. **(a)** Vector genome copy numbers quantified by qPCR in kidneys from mice injected with various AAV vectors via IV injection or unilateral RP injection. Data are presented as double-stranded vector genomes per diploid genomic equivalent (ds-vg/dge). X-axis labels provide information about Groups (*i.e.*, G1, G2, G3, etc.), vectors, and route of administration. G12-LK/G13-LK and G12-RK/G13-RK represent the AAV vector-injected left kidney (LK) and contralateral uninjected right kidney (RK) from Groups 12 and 13, respectively. **(b)** Statistical comparison of vector genome copy numbers between kidneys exhibiting pseudotransduction only (G12-RK, G13-LK, G13-RK) and kidneys with both pseudotransduction and true transduction (G12-LK). Data from G12-RK, G13-LK, G13-RK were combined for robust statistical comparison. **(c)** Transgene mRNA transcript levels quantified by RT-qPCR using the 2^-ΔΔCt^ method with mouse β-actin serving as the internal reference. Data were further normalized to the average of transcript levels in Group 1 for GFP, Group 2 for mCherry and nlsmCherry, and Group 3 for tdTomato, and presented in a graph. **(d)** Statistical comparison of mRNA transcript levels between kidneys with pseudotransduction only and kidneys with both pseudo- and true transduction in PTs. Data are presented as mean ± SD. In Panels b and d, statistical comparisons between the two groups were performed using two-tailed Mann-Whitney U test. Each data point represents an individual kidney. Notably, kidneys with true transduction exhibit significantly higher vector genome copy numbers and transgene transcript levels compared to kidneys with pseudotransduction alone, despite comparable fluorescent reporter protein signals observed by immunofluorescence microscopy (see Fig. 5a **Supplementary** and Fig. 4a),

### Computational quantification reveals binary pseudotransduction versus gradient true transduction in PTs

Having obtained a comprehensive collection of immunofluorescence microscopy images of kidneys exhibiting both pseudotransduction and true transduction in PTs across multiple experimental groups, we were able to systematically examine the spatial distribution patterns of fluorescent reporter signals characteristic of each phenomenon. To objectively assess and quantify both pseudo- and true transductions in PTs, we developed an AI-based computational tool and implemented it as a Python package, kidneyseg_pt. This package automatically segments individual PTs, divides each PT into 50 partitions by uniformly distributed points on each PT outer wall (outer contour), and quantifies the percentage of transduced PTs among all PTs, the percentage of transduction in individual PTs, and the fluorescence signal intensity within each PT in an objective manner (**Fig. 7a, b**). To quantify transduction in PTs, the fluorescent reporter signals were binarized as positive and negative by masking using defined cutoff values, and each partition was scored as positive if any positive reporter signal that meets inclusion criteria was present in its area. The transduction percentage in each PT was then determined as the proportion of positive partitions out of 50 total partitions. Please note that PT cell boundary-based quantification was not possible because individual epithelial cell boundaries were not visible in our immunofluorescence microscope images. In addition, the kidneyseg_pt approach prioritized equal arc-length spacing along the PT outer wall over achieving uniform partition sizes with partitioning lines because the former captures better transduction patterns that follow the epithelial monolayer organization than the latter.

**Fig. 7.**
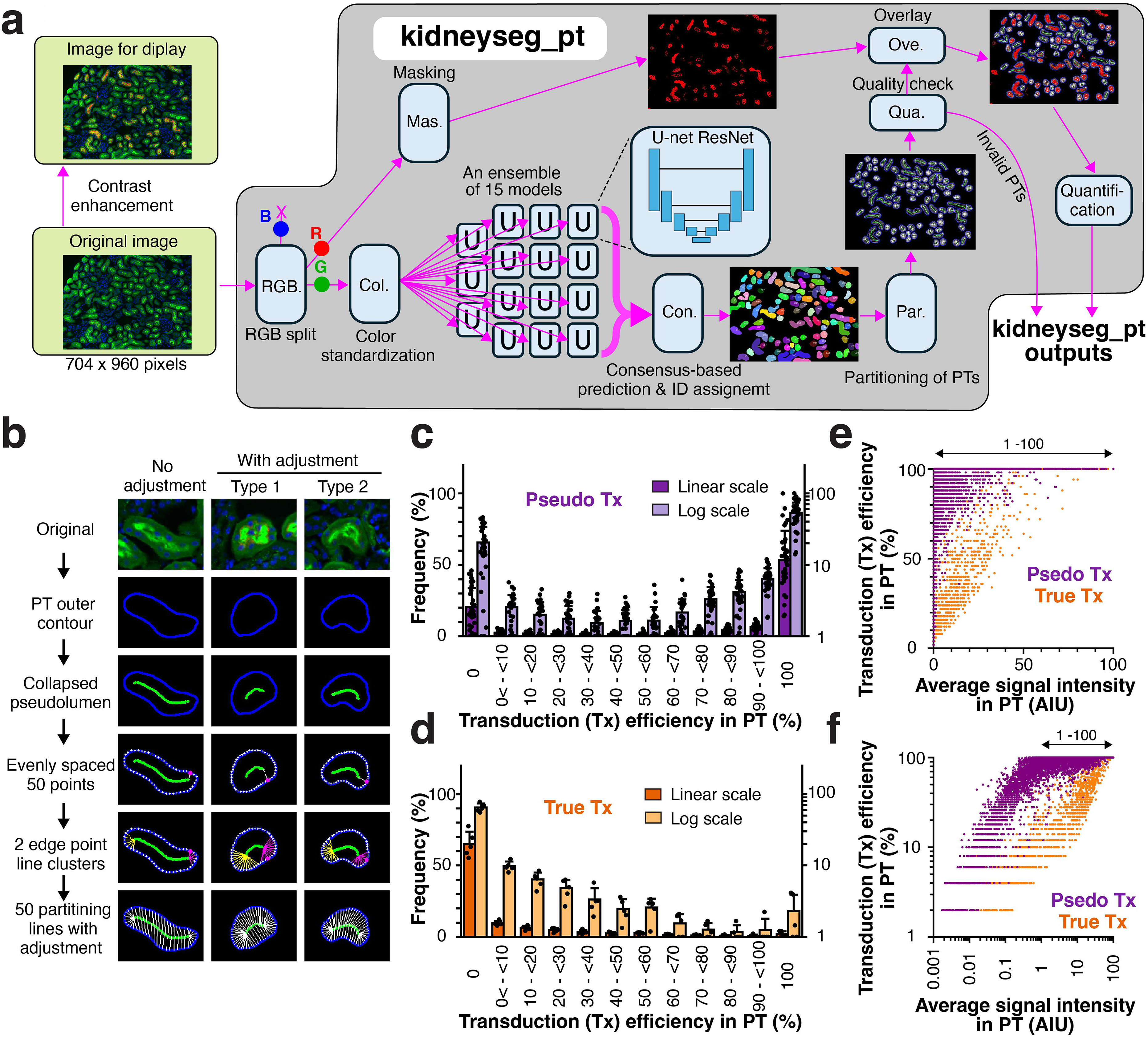
AI-based computational analysis reveals binary pseudotransduction versus gradient true transduction patterns in PTs. **(a)** Workflow diagram of the computational pipeline of the kidneyseg_pt Python package. Original immunofluorescence microscopy images are processed through RGB channel splitting and color standardization. Please note that fluorescent reporter signals may not be visible in the original images shown, as images used for the kidneyseg_pt analysis must maintain a wide dynamic range to preserve quantitative accuracy and may require contrast enhancement for visualization. An ensemble of 15 U-Net with a ResNet50 encoder models performs consensus-based PT segmentation. The pipeline includes signal masking using defined cutoff values, PT partitioning, quality control checks, and quantitative analysis, generating output images and reports. **(b)** Step-by-step illustration of PT partitioning process. From the original image, kidneyseg_pt extracts PT outer contours, draws collapsed pseudolumens, identifies the point on the PT outer contour nearest to either pseudolumen edge (indicated as a magenta point), and generates 50 evenly spaced points from the magenta point along the PT outer contour. For each of the 50 outer contour points, the algorithm determines the nearest point on the pseudolumen. If the nearest point is one of the two pseudolumen edges, these two points are connected with a line, generating an edge point line. The resulting edge point lines form two distinct clusters (indicated with magenta and yellow lines). Then a total of 50 partitioning lines are generated either with no adjustment or with Type 1 or 2 adjustment to maintain appropriate uniformity. **(c, d)** Histogram analysis of transduction efficiency across individual PTs binned into 12 categories. Data are displayed on both linear scale (dark bars, left y-axis) and log scale (light bars, right y-axis). Data collected from a total of 42 kidneys in Groups 2, 5, 6, 8, 10, 12, 15, 16, 18, and 19 (Panel c) and data collected from a total of 5 archived AAV-KP1 or AAV-KP3-CAG-tdTomato RP-injected kidneys (Panel d) were separately pooled and analyzed, showing a characteristic bimodal distribution and a gradient pattern, respectively. Statistical comparison revealed significant differences between pseudotransduction and true transduction in all percentage bins but Bin 8 (q=0.000017 to 0.017, two-tailed Mann-Whitney U-test with Benjamini-Hochberg FDR adjustment). Data are presented as mean ± SD. **(e, f)** Linear-scale (e) and log-scale (f) scatter plots showing the relationship between average fluorescence signal intensity expressed as arbitrary intensity unit (AIU) and transduction efficiency (%) in individual PTs for pseudotransduction (purple, n=7,281 PTs) and true transduction (peach, n=1,235 PTs). Pearson’s r’s were 0.4193 and 0.8544 for pseudo- and true transduction, respectively. When focused only on 1 to 100^th^ percentile signal intensities, the range indicated with double-headed arrows, Pearson’s r for pseudotransduction was reduced to 0.2403. These patterns indicate that pseudotransduction is a binary phenomenon affecting all epithelial cells within a given PT segment regardless of signal intensity, while true transduction is a gradient phenomenon reflecting cumulative outcomes of independently transduced epithelial cells.

The kidneyseg_pt quantitative analysis of transduction patterns was then applied to epifluorescence microscopy kidney images showing pseudo- and true transduction in PTs as well as those showing no transduction as negative controls. The pseudotransduction images were from mice in Groups 2, 5, 6, 8, 10, 12, 15, 16, 18 and 19. Since there were no kidney images showing only true transduction in PTs in the current study, we used archived mouse kidney tissues from our previously published studies that had been transduced via RP injection with either 3.0 × 10^11^ vg of AAV-KP1-CAG-tdTomato (n=3) or 3.0 × 10^11^ vg of AAV-KP3-CAG-tdTomato (n=2)^12, 15^. These archived kidney tissue samples underwent exactly the same immunofluorescence microscopy procedure as that used in the current study groups. Histogram analysis of transduction percentages in individual PTs across 12 bins (a 0% only left-edge boundary bin, 0< to <10%, 10 to <20%, …, and 90 to <100% bins, and a 100% only right-edge boundary bin) revealed striking qualitative differences between pseudotransduction and true transduction patterns in all percentage bins but Bin 8 (q=0.000021 to 0.029, two-tailed Mann-Whitney U-test with Benjamini-Hochberg FDR adjustment, **Fig. 7c, d, Supplementary Figs. 7, 8 and 9**). Pseudotransduction exhibited a bimodal histogram with distinct peaks at 0% and 100%, indicating a near all-or-none pattern in which individual PTs in kidney sections were either completely non-transduced or uniformly transduced, with minimal intermediate states (**Fig. 7c, Supplementary Figs. 7 and 8**, purple). In contrast, true transduction displayed a histogram with peak frequency in the 0% left-edge boundary bin, with each successive bins showing progressively lower PT transduction percentage (**Fig. 7d, Supplementary Fig. 9**, peach). We also investigated the relationship between average fluorescence intensity and percent transduction efficiency in each transduced PT. Within pseudotransduced PTs, pseudotransduction percentage remained consistently high (∼100%) across a wide range of fluorescent reporter signal intensities (**Fig. 7e and f**, purple). Pearson’s correlation analysis of the entire data set (n=7,281 pseudotransduced PTs) revealed a weak positive correlation (Pearson’s r=0.4193, two-tailed p=4.64 × 10^-308^). When 2,251 PTs showing weak signals defined as <1% of the maximum PT signal intensity were excluded (n=5,030), the correlation weakened further (Pearson’s r=0.2403, two-tailed p=5.60 × 10^-67^). In contrast, a strong positive correlation was observed between transduction percentage and signal intensity in truly transduced PTs (n=1,235, Pearson’s r = 0.8544, two-tailed p<5.00 × 10^-324^).

**Fig. 8.**
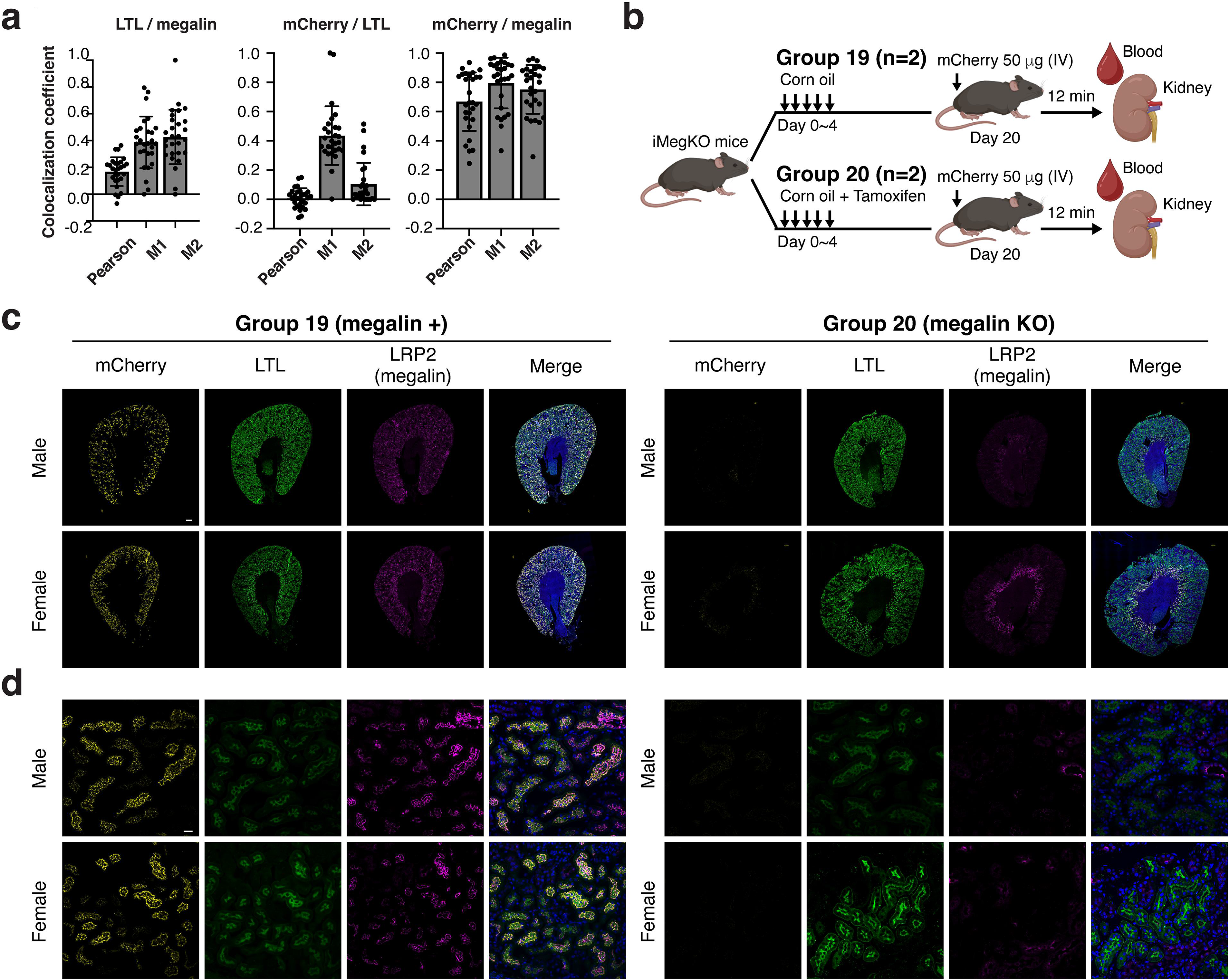
mCherry-mediated pseudotransduction in PTs strongly associates with megalin and is abolished by conditional megalin knockout in mice. **(a)** Quantitative colocalization analysis of LTL, megalin (LRP2), and mCherry signals in kidney sections of AAV vector-treated mice exhibiting pseudotransduction in PTs. To determine Pearson’s correlation coefficient (Pearson), and Manders overlap coefficients (M1 and M2) for LTL/megalin, mCherry/LTL, and mCherry/megalin in each mouse, 2 kidney images acquired with a LSM 900 confocal microscopy using a 20× objective lens were analyzed. Each dot represents a correlation coefficient value obtained from each mouse, and a total of 28 mice from Groups 2, 5, 6, 8, 10, 12, and 13 were analyzed. Data are presented as mean ± SD. **(b)** Experimental design of purified mCherry protein injection in iMegKO mice, a PT-specific inducible megalin knockout mouse model. iMegKO mice (one male and one female per group) received corn oil only (Group 19, megalin+) or corn oil containing tamoxifen (150 mg/kg body weight) for 5 consecutive days (Group 20, megalin KO). On day 20, mice were intravenously injected with 50 μg of purified mCherry protein, and blood and kidneys were collected 12 min post-injection. **(c, d)** Representative epifluorescence images of whole kidney sections (c) and high-magnification confocal microscopy images of the cortex region of kidneys (d) from male and female mice in Group 19 (megalin+, left panels) and Group 20 (megalin KO, right panels). Images show mCherry (yellow), LTL (green), LRP2 (megalin, magenta), and Hoechst 33342 (blue) signals. Control mouse kidneys (Group 19) showed robust megalin expression throughout the kidney and were positive for mCherry signals showing pseudotransduction in PTs. Tamoxifen-treated mice (Group 20) showed loss of megalin expression in the cortex and absence of mCherry pseudotransduction signals in megalin-negative regions in both sexes. Residual megalin expression and its associated mCherry signals were visible in the outer medulla of the female mouse, likely due to incomplete knockout of megalin in this region in females. Scale bars are 300 µm for the whole kidney images and 20 µm for confocal images. High-resolution raw and processed images have been deposited in Zenodo. Image display settings are summarized in **Supplementary Table 3**.

**Fig. 9.**
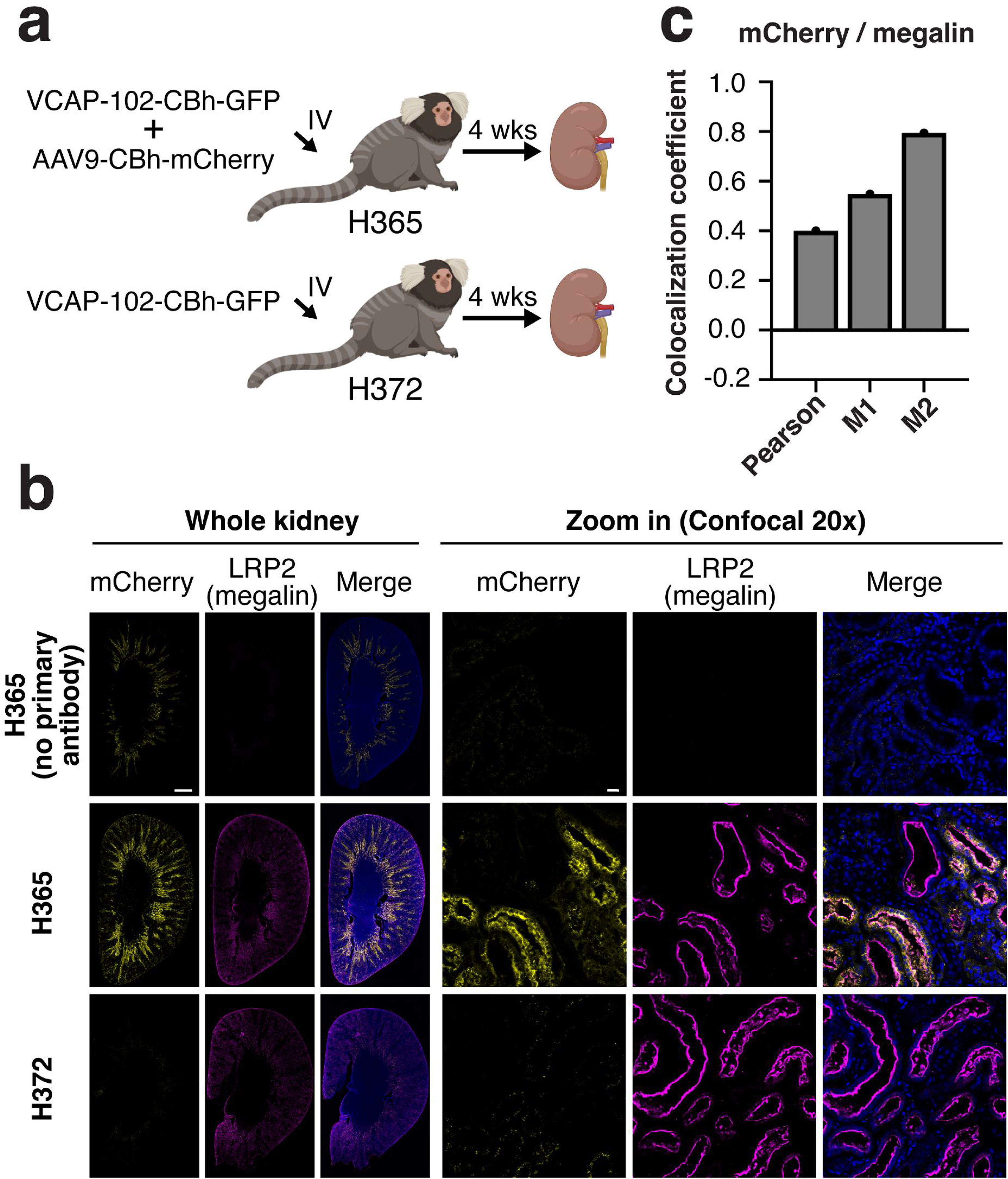
AAV9-mCherry vector-mediated pseudotransduction in PTs in non-human primate. **(a)** Experimental design. Common marmosets H365 and H372 received intravenous injection of AAV vectors at 1.0 × 10^13^ vg/kg per vector and kidneys were collected 4 weeks post-injection. H365 received a 1:1 mixture of AAV9-CBh-mCherry and AAV-VCAP-102-CBh-GFP, while H372 received AAV-VCAP-102-CBh-GFP alone and served as a no-mCherry-vector control. This experimental procedure had been conducted in a previously published study^55^ and the collected kidneys had been fixed with 4% PFA and stored in a refrigerator. These archived kidney samples were subjected to histological assessment in this study. **(b)** Representative whole kidney epifluorescence images (left panels) and high-magnification confocal images (right panels) of kidney sections from marmosets H365 and H372. Secondary antibody only control images without primary antibody against megalin (top row) demonstrated specificity of immunostaining. Kidney sections of H365 immunostained with anti-mCherry and anti-LRP2 (megalin) antibodies (middle row) showed the binary all-or-none punctate luminal and subluminal mCherry signals (yellow) associated with megalin (magenta), consistent with pseudotransduction. Kidney sections from H372 not injected with AAV-mCherry vector showed megalin expression but absence of mCherry signals. Merged images include Hoechst 33342 signals in nuclei (blue). Scale bars are 2 mm for the whole kidney images and 20 µm for confocal images. High-resolution raw and processed images have been deposited in Zenodo. Image display settings are summarized in **Supplementary Table 3**. **(c)** Quantitative colocalization analysis of mCherry and megalin signals in manually segmented mCherry-positive PTs in a confocal kidney section image of H365. Pearson’s correlation coefficient (Pearson), and Manders overlap coefficients (M1 and M2) are shown in a graph.

These observations indicate that pseudotransduction in PT is a binary phenomenon where all epithelial cells within a given PT segment are uniformly affected, resulting in ∼100% pseudotransduction with reporter signal intensities ranging from weak to strong, while other segments remain completely untransduced. In contrast, true transduction is a gradient phenomenon where average signal intensity per area unit in each PT reflects the cumulative outcome of truly transduced epithelial cells, each of which has an independent probability of transduction.

### Megalin-mediated uptake mediates pseudotransduction in PTs

Small circulating proteins, including monomeric fluorescent reporters, pass through the glomerular filtration barrier, are captured by megalin or cubilin, and are reabsorbed into PTs via megalin-mediated endocytosis^35, 36^. Cubilin lacks endocytic capacity on its own and requires cooperation with megalin for internalization^49, 50^. Given this role of megalin and our fluorescence microscopy observations, we hypothesized that megalin plays a central role in mediating pseudotransduction in PTs exposed to GFP or mCherry proteins. To further understand the interaction between megalin and fluorescent reporter proteins, we determined Pearson’s correlation coefficient (PCC) and Manders overlap coefficients (MOCs) between fluorescent reporter signals, megalin signals, and LTL signals using confocal microscopy images of kidney sections in AAV-mCherry vector-injected mice (Groups 2, 5, 6, 8, 10, 12, and 13), which exhibited pseudotransduction with mCherry. PCC measures covariance of two signals but does not directly quantify spatial overlap, while MOCs directly measure spatial overlap between signals above threshold but does not assess intensity relationship. These quantitative analyses revealed that 80% of mCherry signals overlapped with megalin (M1 = 0.80 ± 0.17, means ± SD) and 75% of megalin signals overlapped with mCherry (M2 = 0.75 ± 0.17), indicating substantial spatial coincidence (**Fig. 8a**). However, PCC showed weaker positive correlation (PCC = 0.67 ± 0.20). These observations are consistent with the pattern of pseudotransduction in PTs occurring in nearly all PT epithelial cells with a wide range of fluorescent protein signal intensities (**Fig. 7e and f**, purple). In contrast, mCherry-LTL colocalization was minimal (M1 = 0.44 ± 0.20, and M2 = 0.10 ± 0.14) with no correlation with intensity (PCC = 0.01 ± 0.07) (**Fig. 8a**). PCC and MOC values for LTL-megalin correlation (PCC = 0.17 ± 0.11, M1 = 0.39 ± 0.19, and M2 = 0.43 ± 0.20, **Fig. 8a**) were consistent with the known spatial organization of LTL-binding glycans present only on the surface and the megalin expressed not only on the apical membranes but also in endosomes in PTs. Importantly, all PTs showing pseudotransduction with mCherry expressed megalin at varying levels, and none of pseudotransduced PTs are megalin-negative PTs while there were megalin-positive PTs showing no pseudotransduction (**Fig. 3**, and **Supplementary Fig. 4,** see **Supplementary Discussion 1**).

To show direct evidence that megalin mediates pseudotransduction, we utilized a PT-specific inducible megalin knockout (KO) mouse line, *megalin(lox/lox);Ndrg1-CreER^T^*^2^ (iMegKO)^51, 52^. In this model, tamoxifen (Tam) administration drives Cre-mediated recombination specifically in PT epithelial cells expressing the *N-myc downstream regulated gene 1 (Ndrg1)* gene product, NDRG1, resulting in selective ablation of megalin in PTs in the kidney. This system enabled us to determine whether pseudotransduction is abolished in the absence of megalin expression. Fifty µg of purified mCherry protein was intravenously injected to one male and one female iMegKO mice in which megalin expression in PTs had been conditionally knocked out by intraperitoneal administration of corn oil containing Tam (Group 20) and one male and one female iMegKO mice receiving corn oil only as controls (Group 19). Kidneys and serum samples were then collected 12 min post-injection (**Fig. 8b**). These mice were at ages between 11 to 13-week-old when mCherry protein was injected. In the sera, the presence of circulating mCherry protein was demonstrated in all the 4 mice by the fluorometric assay (**Fig. 2c**). In the kidneys of the control iMegKO mice expressing megalin (Group 19), pseudotransduction in PTs was observed as expected (**Fig. 8c, d**). In both male and female Tam-treated iMegKO mice (Group 20), loss of megalin expression was demonstrated in the cortex (**Fig. 8c, d**). In the outer medulla, megalin expression was lost in the Tam-treated male mouse, but some residual megalin expression was observed in the tam-treated female mouse (**Fig. 8c**). NDRG1 expression can be upregulated by androgen^53^, and sexual dimorphism in gene expression in PTs has been established, with NDRG1 being male biased in S3^54^. Thus, the more complete loss of megalin expression in PTs in the outer medulla corresponding to the S3 segment in the male mouse likely reflects the male-biased NDRG1 expression in S3, resulting in more effective CreER-mediated floxed megalin excision in the male compared to the female mouse. Nonetheless, pseudotransduction was not observed in PTs negative for megalin expression (**Fig. 8d**), demonstrating that megalin plays a crucial role in mediating pseudotransduction in PTs.

### AAV-mCherry vector-mediated pseudotransduction in PTs in non-human primate

Finally, we investigated whether the phenomenon of pseudotransduction in PTs is mouse specific or extends to other animal species. To this end, we analyzed archived common marmoset kidney tissues from a published study conducted at Gumma University^55^. This study systematically compared 11 blood-brain barrier (BBB)-penetrant AAV capsid variants for brain gene delivery following systemic administration in marmosets and identified AAV VCAP-102 as the only capsid outperforming AAV9 in mediating brain-wide transduction; however, kidney transduction remained to be investigated. In the current study, kidneys from the following two marmosets in this previous study were subjected to immunofluorescence microscopy analyses: Animal ID H365 (2.0-year-old female), which received intravenous injection of a 1:1 mixture of AAV9-CBh-mCherry and AAV-VCAP-102-CBh-GFP at 1.0 × 10^13^ vg/kg per vector and Animal ID H372 (2.2-year-old female), which received intravenous injection of AAV-VCAP-102-CBh-GFP at 1.0 × 10^13^ vg/kg (**Fig. 9a**). This design enabled investigation of mCherry-mediated transduction in PTs by comparing H365 (AAV9-CBh-mCherry vector-injected) with H372 (no mCherry vector-injected control). Whole kidney images acquired by epifluorescence microscopy revealed megalin expression throughout the cortex and outer medulla in both H365 and H372, which was absent in H372 kidney sections stained with only secondary antibody, confirming a high signal-to-noise ratio of the megalin signals (**Fig. 9b**). mCherry signals were observed only in H365, detectable as antibody-enhanced signals as well as weaker native fluorescent signals (**Fig. 9b**). Confocal microscopy analysis revealed the characteristic binary all-or-none transduction pattern in PTs indicative of pseudotransduction (**Fig. 9b**), and PCC and MOC assessment demonstrated patterns consistent with mCherry and megalin signal covariance and colocalization in pseudotransduction (**Fig. 9c**). Thus, these findings demonstrate that megalin-mediated pseudotransduction of AAV-mCherry in PTs extends beyond rodents to non-human primates, suggesting that this phenomenon may be broadly conserved across mammalian species.

## Discussion

In this study, we have identified and characterized a previously underappreciated, long-overlooked phenomenon of AAV vector-mediated pseudotransduction in PTs driven by the intrinsic properties of transgene products themselves. This overlooked phenomenon appears to have unknowingly confounded the field of vector-mediated gene delivery to the kidney for decades and significantly contributed to the contradictory observations reported across different studies. The data presented here reconcile apparently discrepant observations in the field, provide new mechanistic insights into how filtered marker gene protein products can confound the interpretation of vector-mediated renal transduction patterns due to pseudotransduction, and underscore the importance of selecting appropriate reporters when evaluating renal gene delivery. This study clearly demonstrates that GFP and mCherry are not suitable for this purpose.

Multiple lines of evidence establish that administration of AAV vectors expressing mCherry or GFP results in pseudotransduction in PTs: (1) Expected nuclear localization in vector-transduced cells was present in reporter protein signals in the liver and heart but was absent in those in PTs despite NLS tagging. (2) We detected circulating mCherry and GFP proteins (<30 kDa) in the blood following AAV vector administration, and previous studies have demonstrated that small proteins of <60 kDa are filtered through the glomerulus and efficiently reabsorbed by PTs via megalin-mediated endocytosis^35, 36, 39^. Therefore, pseudotransduction in PTs with mCherry and GFP proteins represents an expected and scientifically predictable outcome. (3) The observed discordance between readouts of gene delivery (*i.e.*, vector genome copy numbers and transgene transcript levels) and GFP or mCherry reporter protein signals in the kidney contrasts sharply with the concordance in previously demonstrated true transduction^12^. This discordance is also consistent with signals resulting from protein uptake, mediating pseudotransduction. (4) Our experimental approach enabled simultaneous observation of both true transduction and pseudotransduction within the same kidney and the same animal, allowing direct comparison of their distinct histological and molecular signatures using the same vector expressing nlsmCherry (Group 12). Following unilateral RP injection of AAV-KP1-CAG-nlsmCherry, we observed two distinct histological reporter signal patterns within the same injected kidneys. One pattern was consistent with true transduction in PTs characterized by nuclear enriched mCherry signals and another showed all-or-none signals on the luminal surface and in the subluminal cytoplasmic regions consistent with pseudotransduction. In the uninjected kidneys, we only observed the all-or-none pseudotransduction pattern with no nuclear enrichment. In addition, we observed two distinct molecular signatures between RP-injected and uninjected kidneys in the same animals: one consistent with true transduction characterized by concordance of vector genomes, transcripts, and protein expression, and another consistent with protein uptake characterized by discordance. This within-the-same-kidney and within-the-same-animal comparisons in histological and molecular signatures of true- and pseudo-transductions resulting from the same vector provided definitive evidence that supports our conclusion. (5) Immunofluorescence microscopy revealed strong colocalization of fluorescent reporter proteins with megalin on the luminal surface of PTs, along with punctate signals in the subluminal cytoplasmic region that were often not associated with megalin. This subcellular localization of reporter signals is consistent with megalin-mediated endocytosis of reporters and megalin receptor recycling; *i.e.*, initial binding and subsequent clustering on the apical membrane, followed by endocytic internalization, and rapid recycling of megalin receptors back to the cell surface while cargo proteins are sorted for lysosomal degradation^39, 56^. Because mCherry protein demonstrates strong resistance to degradation under the acidic conditions in lysosomes (*i.e.*, pKa <4.5 ^57^), megalin-mediated uptake of circulating mCherry protein leads to its accumulation and retention in endosomal and lysosomal compartments in PTs, generating more prominent pseudotransduction signals compared to other fluorescent proteins with higher pKa values such as GFP (pKa = 6.0). This biochemical distinction of fluorescent reporters elegantly explains why mCherry signals in PTs are consistently reported with AAV-mCherry vectors while GFP signals in PTs exhibit considerable variability across studies^22, 26, 27, 28^. (6) Lastly and importantly, the binary all-or-none transduction pattern of mCherry signals in PTs, regardless of signal intensity, also provides evidence that the observed phenomenon is not a consequence of true AAV vector transduction, which should follow a stochastic model as demonstrated by real-time single AAV particle imaging^31^ and as discussed below.

All biological processes including AAV vector transduction and receptor-mediated reporter protein uptake leading to pseudotransduction are influenced by molecular noise, that is, stochastic fluctuations driven by thermal energy and molecular concentrations^58, 59^. Effects of these random fluctuations are averaged out in large biological systems, such as the kidney as an organ comprising millions to billions of functional units (*i.e.*, cells in the context of transduction) due to the law of large numbers. Consequently, biological outcomes become predictable from given variables such as vector or protein concentration in the system, which follows a deterministic model, although some degree of stochasticity inherently remains at molecular levels. At microscopic levels, such as individual PTs in kidney sections or single PT epithelial cells, stochastic fluctuations can substantially and unpredictably influence biological process outcomes. This stochasticity can be profoundly amplified when biological molecules are present at concentrations below interaction saturation levels or when biological processes involve multiple sequential bottleneck steps as is the case for virus infection and viral vector transduction, where viruses must successfully overcome multi-layer barriers each of which has its own probability of success. In the case of AAV vectors, these barriers comprise receptor binding, endocytosis, endosomal escape, nuclear entry, uncoating, second-strand DNA synthesis, transcription, and translation before achieving productive outcomes. Stochastic failure at any bottleneck step in the AAV vector transduction pathway in cells can prevent productive transduction in an unpredictable manner, resulting in substantial variability in productive outcomes both among individual cells (*e.g.*, PT epithelial cells) and among microscopic functional units (*e.g.*, individual PTs in kidney sections). This results in the generation of mosaic patterns comprising transduced and non-transduced epithelial cells within a PT and varying percentage of transduction among PTs. This phenomenon is exactly what we observed in true transduction in PTs in mice injected with AAV-KP1 vectors via RP. It is important to note that, although all biological processes are inherently stochastic at molecular levels, outcomes could become quasi-deterministic when sequential barriers are only few and individual success probabilities are high. Since pseudotransduction requires only successful receptor binding and endocytosis, which constitute the least restrictive initial steps in cellular uptake, the observed all-or-none pseudotransduction patterns strongly support that pseudotransduction in PTs follows a quasi-deterministic model wherein megalin expression in PTs and its interaction with GFP or mCherry proteins present in urine predetermines outcomes. In this model, all PT epithelial cells expressing megalin take up GFP and mCherry proteins once they appear in the glomerular filtrate in the PT lumens, regardless of protein concentration, resulting in pseudotransduction in all PT epithelial cells with varying signal intensity. This deterministic phenotype contrasts sharply with the mosaic patterns characteristic of true transduction. Interestingly we observed that not all megalin-positive PTs underwent pseudotransduction, revealing an additional layer of binary behavior in pseudotransduction potentially due to internephron heterogeneity^60, 61, 62^ (see **Supplementary Discussion 1**).

Our demonstration of pseudotransduction in PTs and its widespread oversight has significant implications for understanding AAV vector-mediated gene delivery to PTs. This previously unrecognized phenomenon has inadvertently confounded interpretation of experimental data on AAV capsid tropism and transduction efficiency and our understanding of the underlying mechanisms over two decades. A systematic review of published studies reporting AAV-GFP or AAV-mCherry vector-mediated transduction in PTs consistently reveals the binary all-or-none patterns of fluorescent protein signals that are incompatible with the stochastic nature of viral vector transduction and strongly indicate pseudotransduction^21, 22, 23, 26, 27, 28^. Thus, our findings necessitate substantial reassessment of what we have learned from previously published studies and require critical revision of current understanding of AAV-mediated PT transduction as presented in original^21, 22, 23, 26, 27, 28^ and review articles^47, 48, 63, 64^. For example, two high-profile studies have reported substantially superior PT transduction with engineered capsid variants, particularly AAV.cc47, AAV.k13, and AAV.k20, compared to AAV9, following intravenous injection in mice^26, 27^. Interestingly, their studies revealed comparable AAV vector genome copy numbers in the transduced kidneys with AAV9 showing even lower genome copy numbers with statistical significance. They speculated that these engineered capsid-based vectors undergo effective post-entry processes leading to enhanced potency of vector genomes or transcripts. However, all their conclusions on the outcomes of intravenously injected AAV.cc47, AAV.k13, and AAV.k20 vectors were drawn using AAV-CBh-mCherry vectors that effectively transduce the liver and heart. Therefore, we respectfully argue that their conclusions regarding the superior PT transduction with these engineered capsids should be interpreted with extreme caution. Using tdTomato to avoid pseudotransduction in PTs, we could not reproduce their findings and observed no true PT transduction with AAV.cc47, AAV.k13, or AAV.k20 vector following IV injection in mice. In addition, in our head-to-head comparison experiments where we investigated pseudotransduction and true transduction in PTs simultaneously within the same kidneys and within the same animals using the same vector, we observed concordance across vector genome copy numbers, vector transcript levels, and protein levels only in true transduction, and pseudotransduction exhibited their discordance as found in their studies. More recently, researchers systematically investigated optimal combinations of 7 parameters including capsid, promoter, fluorescent protein reporter, dose and route of administration to maximize AAV vector-mediated transduction in PTs in mice^28^. They reported that IV administration of AAV9-CBh-mCherry with Woodchuck hepatitis virus posttranscriptional regulatory element (WPRE) achieved as high as 40 to 60% transduction in PTs. Although we acknowledge the rigor of their experimental design and cutting-edge approaches, we respectfully argue that their fundamental assumption that mCherry signal represents bona fide transduction is likely incorrect as demonstrated in the present study. Consequently, even carefully optimized parameters determined by AAV-mCherry vectors may not reliably enhance true transduction efficiency in PTs. In their study, the authors performed a head-to-head comparison of fluorescent reporter signal intensity between AAV9-CBh-mCherry-WPRE and AAV9-CBh-GFP-WPRE following IV injection in mice, including a co-injection experiment. They demonstrated that AAV9-CBh-mCherry-WPRE yielded robust mCherry signals, whereas GFP signals were barely detectable in the kidneys. This observation is consistent with our inference that mCherry exhibits greater resistance to the acidic endosomal and lysosomal environment compared to GFP, resulting in stronger and more persistent pseudotransduction signals in PTs following endocytic uptake. Our study also revealed additional complexities in understanding AAV vector transduction in the kidney that challenge conventional assumptions about vector design modularity. We observed that the choice of fluorescent reporter protein profoundly influenced transduction outcomes in a cell type-specific manner in the kidney, even when capsid and promoter remained the same (see **Supplementary Discussion 2**).

In conclusion, our study resolves a long-standing enigma in AAV vector-mediated gene delivery to the kidney by clearly delineating pseudotransduction versus true transduction in PTs and elucidating the megalin-dependent mechanism underlying pseudotransduction. This enables us to reconcile over two decades of contradictory observations and reveals a critical gap in our understanding of renal gene delivery. Our findings not only establish a rigorous standard for evaluating kidney transduction using appropriate reporters but also underscore the importance of a fundamental reassessment of previous reports.

## Methods

### Cells

Human embryonic kidney (HEK) 293 cells (240073, Agilent, Santa Clara, CA) were maintained in Dulbecco’s Modified Eagle Medium (DMEM) supplemented with 10% (v/v) fetal bovine serum (FBS), 100 U/mL penicillin, 100 µg/mL streptomycin, and 2 mM L-glutamine (*i.e.*, complete medium) at 37°C in a humidified incubator with 5% CO_2_ and passaged every 2 or 3 days using 0.05% trypsin-EDTA.

### AAV vectors and plasmids

AAV9 and AAV-KP1 helper plasmids were provided by J. M. Wilson and M. A. Kay, respectively. pAAV-CAG-tdTomato and pAAV-CAG-nlsGFP plasmids were gifts from E. Boyden and V. Gradinaru (59462 and 104061, Addgene, Watertown, MA), respectively. The AAV helper plasmids for the AAV.cc47, AAV.k13, and AAV.k20 capsids were constructed using the AAV9 helper plasmid as the backbone. Briefly, the DNA sequence of the AAV9 helper plasmid coding amino acid positions 452-458 in the capsid variable region IV (VR-IV) was substituted with the following amino acid-coding DNA sequence: GGTGTGTCTCTCGGTGGTGGC coding GVSLGG for AAV.cc47, ATGGTGGAAGGACGAATGGGT coding MVEGRMG for AAV.k13, and GTGTCTGGACGTCGCGAAGGT coding VSGRREG for AAV.k20. To construct pAAV-CAG-mCherry, the open reading frame (ORF) of tdTomato was replaced with that of mCherry. To construct pAAV-CAG-nlsmCherry, the NLS coding sequence from pAAV-CAG-nlsGFP was added to the 5’ end of the mCherry ORF in pAAV-CAG-mCherry. A liver-specific promoter (LSP) was designed by fusing three elements in the following order from 5’ to 3’: the hepatocyte-specific cis-regulatory module 8 (HS-CRM8)^65^, the mouse transthyretin gene promoter region (chr18:20,664,319-20,666,325; mm10/GRCm38 assembly), and a 92-bp intron-containing sequence from the murine virus of mice (MVM) genome (positions 2312-2400; GenBank accession NC_001510.1) and synthesized by GenScript Biotech Corporation. To construct pAAV-LSP-nlsmCherry, the CAG promoter in pAAV-CAG-nlsmCherry was replaced with the LSP promoter. All the plasmids were grown in ElectroMAX DH10B cells (18290015, Thermo Fisher Scientific, Hampton, NH), and their sequences were validated by whole plasmid sequencing conducted by Plasmidsaurus Inc. All the AAV vectors used in the study were produced in HEK293 cells by an adenovirus-free plasmid transfection method and purified by two rounds of cesium chloride density-gradient ultracentrifugation, followed by dialysis. AAV vector titers were determined as described previously^66, 67^.

### Mouse experiments

All mouse experiments were approved by the Institutional Animal Care and Use Committee of Oregon Health & Science University (Approval No. TR02_IP00001195) and performed according to the guidelines for animal care at the university. Mice were kept with free access to food (5L0D, LabDiet, St. Louis, MO) and water in a controlled environment (12 h light-dark cycle with an average humidity range of 30-70% and temperatures of 20-23 °C). Eight-week-old C57BL/6J (Strain ID: 000664) male mice were purchased from the Jackson Laboratory. The proximal tubule-specific inducible megalin knockout (KO) mouse line, *megalin(lox/lox);Ndrg1-CreER^T^*^2^ (iMegKO)^51, 52^ was generously provided by Kiyoshi Mori, Kumamoto University, Japan). Mice were genotyped by quantitative PCR (Transnetyx, Cordova, TN) prior to experimentation. Conditional deletion of megalin in PTs was induced by intraperitoneal administration of tamoxifen (Tam) in corn oil (150 mg/kg body weight) once daily for 5 consecutive days. To prepare Tam in corn oil, 20 mL of corn oil (C-8267, Sigma-Aldrich, St Louis, MO) was heated at 42 °C for 1.5 h, 0.2 g of Tam was added to the heated corn oil, and the oil was stirred at 37 °C for 3 h. Both corn oil only and corn oil containing Tam were sterilized using 0.22 µm Centrifuge Tube Top Vacuum Cellulose Acetate Membrane Filter (09-761-34, Corning, Tewksbury, MA). Test agents (purified mCherry protein in this study) were administered into iMegKO 15 days after the completion of the corn oil treatment (corn oil only and corn oil containing Tam). Both male and female mice were used for the study.

Eight-week-old C57BL/6J male mice were injected with AAV vectors (AAV9, AAV.cc47, AAV.k13, and AAV.k20 vectors), AAV vector plasmids (pAAV-CAG-nlsmCherry and pAAV-LSP-nlsmCherry), mCherry protein, or saline. Untreated age-matched C57BL/6J male mice were also included as controls. **Table 1** provides a detailed summary of the experimental groups. In brief, for the AAV9, AAV.cc47, AAV.k13 and AAV.k20 capsid validation studies, mice were injected with AAV vectors via the tail vein (*i.e.*, IV injection) at 4 × 10^13^ vg/kg or 3 × 10^11^ vg/mouse (*n* = 3 to 6 mice per group) or via unilateral renal pelvis (RP) injection into the left kidney at 3 × 10^11^ vg/mouse (n=2 to 3 mice per group). These procedures have been described previously^12, 15^. Two weeks or 30 days post-injection, mice were euthanized, and kidneys, livers and hearts were harvested as described below for histological assessment of vector transduction and molecular analyses. For the hydrodynamic tail vein injection^42^, 100 mL/kg saline containing 50 µg of plasmid DNA or saline only was rapidly injected via the tail vein within 5 sec. Three days or 7 days post-injection, mice were euthanized, and kidneys were harvested as described below for histological assessment. For the mCherry protein injection, 8-week-old C57BL/6J male mouse (n=1), 11-week-old Tam-containing corn oil-treated iMegKO male and female mice (n=1 per sex), and 13-week-old control corn oil-treated iMegKO male and female mice (n=1 per sex) were injected with 50 µg of purified mCherry protein with N-terminal HIS tag, expressed in E. coli (TP790040, OriGene Technologies, Rockville, MD) via the tail vein. Twelve minutes post-injection, mice were euthanized, and kidneys were harvested for histological assessment as described below. Serum samples were collected at the study endpoints in all the mice as well as before agent administration in mice in Group 10, and stored at -80 °C until use. At euthanasia, mice were deeply anesthetized with isoflurane and transcardially perfused with 20 mL of ice-cold phosphate-buffered saline (PBS) to clear the vasculature. Harvested tissues were immediately partitioned, and either post-fixed in ice-cold 4% paraformaldehyde (PFA) in phosphate buffer (PB) for histological analyses or snap-frozen in liquid nitrogen and stored at −80 °C for molecular analyses.

### Archived marmoset kidney samples for histological analysis

Archived kidney samples from two common marmosets (*Callithrix jacchus*) were used in this study. These samples were originally obtained from animals housed at the Gunma University Bioresource Center as part of a previously published AAV gene delivery study^55^. Briefly, animals received intravenous injection of AAV9-CBh-mCherry or a mixture of AAV9-CBh-mCherry and AAV-VCAP-102-CBh-mGFP. Four weeks post-injection, kidney tissues were harvested following in vivo perfusion with ice-cold D-PBS (–) (14249-95, Nacalai Tesque, Kyoto, Japan) containing 20 mM EDTA (311-90075, Nippon Gene, Tokyo, Japan) followed by 4% PFA in 0.1 M PB (006775-1L, Bioenno Lifesciences, Santa Ana, CA). Tissues were then immersed in 4% PFA in PB overnight at 4 °C, washed in PBS (T9181, Takara Bio, Kusatsu, Japan), and stored in PBS supplemented with 0.03% sodium azide at 4 °C until use. All original animal procedures were approved by the Institutional Animal Care and Use Committee of Gunma University (approval Nos. 18-019, 20-053, 21-063, 23-018, 23-057, and 24-057) and complied with the Guide for the Care and Use of Laboratory Animals and the Guidelines for Experiments on Primates in the Field of Neuroscience by the Japan Neuroscience Society.

### AAV vector genome DNA copy number quantification

Total DNA was extracted from tissues using the QIAmp DNA Mini Kit (51304, Qiagen, Venlo, Netherlands) according to the manufacturer’s protocol and eluted in 100 µL of 1 × Tris-EDTA (TE) buffer (pH 8.0). AAV vector genome copy numbers in samples were then quantified by quantitative PCR (qPCR). In brief, 100 ng of tissue DNA was mixed with 12.5 µL of Power SYBR Green PCR Master Mix (4367659, Thermo Fisher Scientific) and 25 pmol primers in a 25 µL reaction volume and subjected to qPCR using Rotor-Gene Q (Qiagen). We amplified the nlsGFP, mCherry/nlsmCherry and tdTomato gene sequences using gene specific primers (**Supplementary Table 1**). qPCR standards were created using pAAV-CAG-nlsGFP, pAAV-CAG-tdTomato, or pAAV-CAG-mCherry. Further details regarding copy number standards and normalization using the *Agouti* gene as an endogenous control were described previously^68^. Vector genome copy numbers are expressed as double-stranded vector genome copy numbers per diploid genomic equivalent (ds-vg/dge). The qPCR amplification conditions were as follows: 95 °C for 10 min, followed by 40 cycles of 95 °C for 15 sec and 60 °C for 1 min.

### Quantification of AAV vector genome transcripts by reverse transcription (RT)-PCR

Total RNA was extracted from tissues using TRIzol Reagent (15596018, Thermo Fisher Scientific). RNA was dissolved in 25 µL of nuclease-free water (HyClone HyPure Water, Molecular Biology Grade, SH30538.03, Cytiva, Marlborough, MA). Two µg of total RNA was treated with DNase (TURBO DNA-free Kit, AM1907, Thermo Fisher Scientific) in a 21 µL reaction mixture for 1 h at 37 °C. Eleven µL of this mixture containing approximately 1 µg of DNase-treated RNA was then subjected to complementary DNA (cDNA) synthesis using the SuperScript IV First-Strand Synthesis System (18091200, Thermo Fisher Scientific) with an oligo(dT)_20_ primer in a 20 µL reaction mixture according to the manufacturer’s instruction. After the completion of cDNA synthesis, the mixture was made to 100 µL by adding 80 µL of the nuclease-free water. For RT-qPCR, 5 µL of the diluted cDNA was mixed with 12.5 µL of the Power SYBR Green PCR Master Mix and 25 pmol of each primer (a pair of primers to amply the target transgene transcripts and another pair for β-actin transcripts) in a 25 µL reaction volume and subjected to qPCR. Relative mRNA expression levels of target transgenes were calculated using the 2^-ΔΔCt^ method with mouse β-actin serving as the internal reference. Data were further normalized to the average of transcript levels in Group 1 for GFP, Group 2 for mCherry and nlsmCherry, and Group 3 for tdTomato. All qPCR primer sequences are detailed in **Supplementary Table 1**.

### Histological processing and analysis

Organs (kidneys, livers, and hearts) were harvested from mice following perfusion with ice-cold PBS and fixed in ice-cold 4% PFA (158127, Sigma-Aldrich) overnight. PFA-fixed organs were then equilibrated in 30% sucrose (S5, Thermo Fisher Scientific) for cryoprotection. Subsequently, the fixed kidney and heart tissues were placed in 15 mm × 15 mm cryomolds (25608-924, Avantor, Radnor Township, PA) and Tissue-Tek O.C.T. Compound (25608-930, Avantor) was poured over the tissue. The fixed tissue-containing molds were then flash-frozen in a dry ice-2-methylbutane (MX0760-1, MilliporeSigma, Burlington, MA) bath and stored at −80 °C until use. Liver tissues were embedded in 4% agarose (Agarose RA, N605-500G, Avantor). In brief, a 4% agarose solution was prepared by dissolving 4 g agarose in 100 mL of PBS using a microwave. Liver samples were placed in a mold (25608-916, Avantor), and molten agarose was poured over the tissue and allowed to solidify at room temperature for 10 min. The embedded tissue block was mounted on a Leica vibrating blade microtome (VT1000S, Leica Biosystems Nussloch, GmbH, Germany) and sectioned in PBS at a blade advance speed setting of 9. Floating sections (60 µm) were collected, transferred to 24-well plates (353047, Corning, Glendale, AZ) containing PBS supplemented with 0.02% sodium azide (S2002, MilliporeSigma), and stored at 4 °C until use.

For immunofluorescent microscopy, frozen kidney and heart tissues were sectioned at 5-µm thickness using a cryostat (TN50, Tanner Scientific, Bradenton, FL). Kidney sections were blocked with 1% bovine serum albumin (BSA, A3294, Sigma-Aldrich) in PBS at room temperature for 1 h, followed by incubation with primary affinity reagents diluted in 1% BSA at 4 °C overnight. The primary affinity reagents used in this study were: rabbit anti-WT1 (1:100, 83535, Cell Signaling Technology, Danvers, MA), rabbit anti-LRP2 (1:400, ab76969, Abcam, Waltham, MA), and LTL-fluorescein (1:200, FL-1321-2, Vector Laboratories, Newark, CA). Sections were then incubated with Alexa Fluor 647-conjugated goat anti-rabbit IgG (Alexa Fluor 647 AffiniPure Goat Anti-Rabbit IgG (H+L), 1:500, 111-605-144, Jackson ImmunoResearch, West Grove, PA) in 1% BSA at room temperature for 1 h. Following secondary antibody incubation, slides were washed in PBS for 10 min and stained with Hoechst 33342 (1:4,000,000, H3570, Thermo Fisher Scientific) for 10 min, followed by two PBS washes (3 min each). Heart sections were stained with Hoechst only. For floating liver sections, samples were washed in PBS at room temperature for 10 min, stained with Hoechst (1 µg/mL) for 30 min, and washed again in PBS for 10 min. After the staining was completed, a small amount of Fluoromount-G (0100-01, SouthernBiotech, Birmingham, AL, USA) was dropped near the sample, and a coverslip (12541031, Thermo Fisher Scientific) was placed on top of the sample. Immunofluorescence images were acquired using a Keyence BZ-X710 (Keyence, Osaka, Japan) for epifluorescence microscopy and an LSM 900 with Airyscan 2 (Carl Zeiss, Oberkochen, Germany) for confocal microscopy.

Fluorescence images of mouse tissues were acquired in the following manner. For whole organ imaging of the kidney and the heart, images of individual fields of view were acquired using a Keyence BZ-X710 (Keyence, Osaka, Japan) with a 20× objective lens. Acquired individual image tiles were then stitched using Keyence BZ-X Analyzer software (version 1.4.1.1) to generate fully reconstructed, composite whole organ images. Whole liver images were acquired under the widefield setting using an LSM 900 confocal laser scanning microscope equipped with Airyscan 2 and a 10× objective lens. The acquired liver image tiles were subsequently stitched using ZEN software (Carl Zeiss) with default parameters, utilizing the DAPI channel as the reference channel for alignment and fusion. Exposure times for epifluorescence images and parameters for confocal images were held constant for each specific channel across all comparable samples.

For high-magnification epifluorescence microscopy imaging of the kidney, images were acquired using BZ-X710 with a 20× objective lens. Exposure times were held constant for each specific channel across all comparable samples. For the mCherry channel to detect signals from kidney samples, dual exposure times of 20 ms and 3 ms were utilized. The 20-ms exposure was used to generate representative images for visual display in figures (**Supplementary Fig. 4a**), whereas the 3-ms exposure was used to prevent signal saturation and preserve a wide dynamic range for computational quantification of PT transduction using the kidneyseg_pt Python package. For the mCherry signal detection for the kidneyseg_pt-based quantification analysis, we selected 3 ms exposure time capable of capturing both weak signals (*e.g.*, Groups 18 and 19) unambiguously and strong signals (*e.g.*, Groups 6 and 16) without significant pixel saturation using the same image acquisition parameters. Confocal microscopy of kidney and liver tissues was performed using an LSM 900 confocal laser scanning microscope with 10× and 20× objective lenses. Image acquisition conditions for each selected dye were automatically configured using the Zen software’s Smart Setup wizard to maximize signal gain while minimizing crosstalk. Detector gain settings were maintained between 550 and 850 V, and laser transmission percentage was adjusted between 0.1% and 5% depending on the fluorophore and tissue sample. The pinhole size was set to 1.4 Airy Units (AU) for the Hoechst 33342 channel and 1.0 AU for all other fluorophore channels. Digital offset was set to 0, and digital gain was set to 1.0 for all acquisitions. Data were acquired at 16-bit depth per channel using the ZEN software’s Smartest (sequential line scanning) mode with 2x averaging. Acquisition parameters were kept constant for each specific channel across all comparable samples. For all subsequent visual presentation and quantitative analyses, images were processed and exported at an 8-bit depth per channel. Liver confocal images were acquired primarily in the same manner. Please note that mCherry signals in epifluorescence whole kidney images (**Fig. 8**) were acquired using a longer exposure time (400 ms) compared to other experimental groups (50 ms in **Fig. 3**). Similarly, mCherry signals in high-magnification confocal kidney images (**Fig. 8**) were acquired using a higher laser transmission percentage (2%) compared to the standard mCherry condition used for other groups (0.2% in **Fig. 3**). Because mCherry signals in kidneys from iMegKO mice receiving purified mCherry protein (Groups 19 and 20) were substantially weaker than those from other experimental groups, these enhanced acquisition parameters were necessary to improve signal-to-noise ratio and enable clear visualization of these weaker signals without excessive adjustment of image display parameters.

Immunofluorescence microscopy of 4% PFA-fixed, archived marmoset kidney tissues stored at 4 °C was performed in the following manner. Kidneys were bisected with a scalpel and the half-kidney was embedded in 2% agarose gel in PBS. One hundred-μm-thick sections were obtained using a vibrating blade microtome (VT1200S, Leica) and stored at 4°C in PBS containing sodium azide until use. Tissue sections were incubated overnight at room temperature with the following primary antibodies diluted in blocking solution (2% donkey serum [S30-100ML, Merck, Darmstadt, Germany], BSA [A9647, Merck], 0.5% Triton X-100, and 0.03% sodium azide in PBS): rat monoclonal anti-mCherry antibody (1:500, M11217, Thermo Fisher Scientific), rabbit polyclonal anti-LRP2 antibody (1:100, ab76969, Abcam). For visualization, sections were incubated at room temperature for 3 to 4 h in blocking solution containing the following secondary antibodies and Hoechst 33342 (1:1,000, R37605, Thermo Fisher Scientific): donkey anti-rat IgG Alexa Fluor Plus 555 (1:2,000, A48270, Thermo Fisher Scientific) and donkey anti-rabbit IgG Alexa Fluor Plus 647 (1:2,000, A31573, Thermo Fisher Scientific). Following secondary antibody incubation, sections were mounted on glass slides using ProLong Glass Antifade Mountant (P36980, Thermo Fisher Scientific), allowed to cure at room temperature for 3 to 7 days, and stored at 4 °C. For whole organ imaging of the kidney, images of individual fields of view were acquired using a Keyence BZ-X800 (Keyence, Osaka, Japan) with a 2× objective lens with constant exposure times adjusted for each channel. Acquired individual image tiles were processed as described to generate whole organ images. Confocal microscopy was performed using an LSM 800 confocal laser scanning microscope (Carl Zeiss) with a 20× objective lens. Image acquisition parameters were configured for multi-channel image acquisition in the same manner as described above. The pinhole size was set to 1.12 AU for the nuclear channel, 0.92 AU for the mCherry channel, and 0.78 AU for the megalin channel. Detector gain settings were maintained between 670 and 850 V, digital gain was adjusted between 1.0 and 3.0, and digital offset ranged from -20000 to -15000 depending on the fluorophore and tissue sample. Laser transmission percentage was adjusted to 58%. Data were acquired at 16-bit depth per channel using sequential line-scanning mode with 2× frame averaging. Acquisition parameters were kept constant for each specific channel. For all subsequent visual presentation and quantitative analyses, images were processed and exported at 8-bit depth per channel.

### Post-processing of fluorescence microscope images for visual display

Background fluorescence thresholds for green, red, and far-red channels were determined using the 99^th^ percentile method from negative control samples, including tissue sections from untreated animals or sections stained with secondary antibody only (no primary antibody). Background thresholds were applied as minimum display values. Maximum display values were set to 255 as default for 8-bit images unless contrast enhancement was required for clear visualization of signals in figure images. No gamma correction was applied. Image display parameters were held constant for fluorescent protein marker channels across all comparable samples. Parameters for other channels were also held constant with some exceptions for clear presentation. Image display settings used for figures are fully disclosed in **Supplementary Table 3**. Raw image files at an 8-bit depth per channel 8-bit have been deposited in Zenodo.

### Quantitative colocalization analysis of confocal images of kidney sections

Colocalization of dual-channel signal combinations, LTL versus megalin, mCherry versus LTL, and mCherry versus megalin, was quantified using Pearson’s correlation coefficient (PCC) and Manders overlap coefficients (MOCs). mCherry signal-positive PTs were manually segmented using the region of interest (ROI) Manager in ImageJ2 (Fiji), and binary masks were generated from the defined ROIs. Colocalization analysis was then performed exclusively within these ROIs using the Coloc2 plugin in ImageJ2 (Fiji). To exclude background noise objectively, channel-specific intensity thresholds were automatically calculated using Costes threshold regression and used as cutoff values to determine M1 and M2. Using this method, confocal kidney images were assessed in 28 mice across 7 experimental cohorts (3, 3, 3, 3, 6, 4, and 6 mice in Groups 2, 5, 6, 8, 10, 12, and 13, respectively) for **Fig. 8a**, and in one marmoset (H365) for **Fig. 9c**.

### Preparation of HEK293 cell lysates containing fluorescent proteins

HEK293 cell lysates containing nlsGFP, mCherry, nlsmCherry, and tdTomato were generated for use as positive controls in western blot analysis of mouse serum samples. In brief, 4 × 10^6^ HEK293 cells were seeded in 55 cm^2^ Tissue Culture (TC)-Treated Culture Dishes (08-772-22, Fisher Scientific). The following day, complete culture medium was replaced with serum-free DMEM supplemented with penicillin, streptomycin, and L-glutamine, and cells were transfected with 2 µg of plasmid DNA (pAAV-CAG-tdTomato, pAAV-CAG-mCherry, pAAV-CAG-nlsmCherry, or pAAV-CAG-nlsGFP) using our standard polyethylenimine (PEI) transfection method^66^. Two days post-transfection, cells and media were collected, centrifuged at 1,000 × g for 10 min to obtain cell pellets. Cell pellets were washed once with PBS and spun down at 1,000 × g for 5 min. Cell pellets were then lysed in 500 μL of Mammalian Protein Extraction Reagent (78503, Thermo Fisher Scientific) supplemented with Halt Protease Inhibitor (78440, Thermo Fisher Scientific) at a 100:1 (v/v) ratio. After 15-min incubation, lysates were clarified by centrifugation at 16,000 × g at 4°C for 10 min. Protein concentrations were determined using the Pierce BCA Protein Assay Kit (23227, Thermo Fisher Scientific) according to the manufacturer’s instruction. Control HEK293 lysates containing no fluorescent proteins were generated using non-transfected cells in the same manner.

### Western blot analysis of serum samples

Serum samples were diluted at 1:2.5 in PBS by adding 4µL of each serum to 6 µL of PBS, and mixed with 2 µL of 6× sample buffer (375 mM Tris-HCl, 9% SDS, 9% β-mercaptoethanol, 50% glycerol, 0.03% bromophenol blue, pH 6.8). HEK293 cell lysates were diluted with PBS to reach predetermined concentrations (80 µg/mL for the mCherry, nlsmCherry, and tdTomato-containing protein lysates; and 20 µg/mL for the GFP-containing protein lysates), and 10 µL of each diluted lysate was mixed with 2 µL of 6× sample buffer. These samples were then heated at 100 °C for 10 min, loaded onto 7.5% SDS-polyacrylamide mini gels or 8-16% Mini-PROTEAN® TGX™ Precast Protein Gels (4561103, Bio-Rad, Hercules, CA), and electrophoresed at 12 mA in 1× premixed electrophoresis buffer prepared from 10x Tris/Glycine/SDS (25 mM Tris, 192 mM glycine, 0.1% SDS, pH 8.3; 1610732, Bio-Rad) along with 7 µL of Precision Plus Protein Dual Color Standards (1610374, Bio-Rad) for approximately 2 h. Separated proteins in gels were then transferred to polyvinylidene difluoride (PVDF) membranes (IPVH00010, Thermo Fisher Scientific) in Pierce^TM^ 1-StepTransfer Buffer (84731, Thermo Fisher Scientific) using Pierce^TM^ G2 Fast Blotter (62288, Thermo Fisher Scientific) with the mixed MW range setting (25 V and 1.3 A for 7 min). Following transfer, membranes were washed with Milli-Q water for 2 min and then with 1× Tris-buffered saline (TBS) with 0.1% Tween 20 (p7949, Sigma-Aldrich) (TBS-T) for 5 min, followed by blocking with 5% bovine serum albumin (BSA, for mCherry and tdTomato detection) or 5% non-fat dry milk (for GFP detection) prepared in 1× TBS-T at room temperature for 1 h. Membranes were then incubated with respective primary antibodies: HRP-conjugated anti-mCherry recombinant rat monoclonal antibody (1:1000, 740003THRP, Thermo Fisher Scientific) or anti-GFP mouse monoclonal antibody (1:1000, sc-9996, Santa Cruz Biotechnology, Dallas, TX, USA) at 4 °C for approximately 18 h. After completion of primary antibody incubation, membranes were washed three times with 1× TBS-T for 10 min each. Membranes probed with anti-GFP antibody were subsequently incubated with an HRP-conjugated rat monoclonal anti-mouse immunoglobulin antibody, Mouse TrueBlot ULTRA, (1:5000, 18-8817-30, Rockland, Limerick, PA, USA) at room temperature for 1 h followed by 3 washes with 1× TBS-T for 10 min each. Membranes probed with HRP-conjugated anti-mCherry antibody were only washed three times with 1× TBS-T for 10 min each. To detect antibody-bound proteins, anti-mCherry antibody-probed and anti-GFP antibody-probed blots were incubated with SuperSignal^®^ West Femto Luminol/Enhancer Solution (34094, Thermo Fisher Scientific) and MilliporeSigma™ Immobilon™ Western Chemiluminescent HRP Substrate (ECL) (WBKLS0500, Thermo Fisher Scientific) according to the manufacturer’s recommendation, respectively, and signals were captured using Amersham™ ImageQuant™ 800 (29399481, Cytiva). Images were recorded as 10 serial exposures at a 40-s interval.

In one experiment, albumin-depleted sera were prepared from sera collected from mice in Group 10 (AAV.k20-CAG-mCherry, IV) using Albumin Depletion Kit (ab241023, Abcam) according to the manufacturer’s protocol with a minor modification in which we mixed 10 µL of serum and 50 µL of Albumin Binding Buffer provided by the kit. Both serum samples collected before AAV vector injection (negative controls) and those collected 30 days post-injection were subjected to the albumin depletion procedure, and analyzed for the presence of mCherry protein in sera by western blot. Albumin was depleted from samples to reduce background interference and improve band resolution. Western blot using non-depleted serum samples resulted in heavy smearing and high background due to albumin overloading although not preventing mCherry detection. Fifteen µL of each diluted, albumin-depleted sample was mixed with 3 µL of 6× sample buffer and loaded on an 8-16% Mini-PROTEAN^®^ TGX™ Precast Protein Gels together with a non-depleted naive mouse serum and 12 µL of mCherry-containing HEK293 cell lysate described above, and subjected to the above-described western blot analysis probed with anti-mCherry antibody.

### Fluorometric quantification of circulating fluorescent proteins

To quantitatively measure fluorescent proteins circulating in mouse sera, we developed a direct fluorescent protein quantification assay for mCherry, GFP, and tdTomato present in mouse sera. To this end, commercially available purified recombinant proteins, purified mCherry (TP790040, OriGene), purified GFP (GFP, His-Tag Recombinant, 102418-1, BPS Bioscience, San Diego, CA, USA), and purified tdTomato (Purified fluorescent protein tdTomato, with N-terminal HIS tag, expressed in E. coli, TP790045, AMSBIO, Cambridge, MA, USA), were purchased and used as standards. To generate standard curves and determine dynamic range of the assays, purified fluorescent protein standards were serially diluted in the serum-PBS diluent explained below to create 14-point standard curves ranging from 100 µg/mL to 0.01 µg/mL (Standards A to N). The serum-PBS diluent was prepared by diluting naive mouse serum pooled from two untreated 8-week-old C57BL/6J male mice with PBS at 1:7.5 (1 part serum in 6.5 parts of PBS). This dilution matched experimental samples to ensure equivalent serum background across standards and samples. In brief, to create GFP Standard A (100 µg/mL), 14.1 µL of Purified GFP (0.71 mg/mL) was mixed with 85.9 µL of the serum-PBS diluent; to create mCherry Standard A (100 µg/mL), 12.7 µL of Purified mCherry (0.79 mg/mL) was mixed with 87.3 µL of the serum-PBS diluent; and to create tdTomato Standard A (100 µg/mL), 8.0 µL of purified tdTomato (1.25 mg/mL) was mixed with 92.0 µL of the serum-PBS diluent. Subsequent standards were prepared by sequential 1:2 dilutions, mixing 50 µL of the preceding standard with 50 µL of the serum-PBS diluent. Standards A to N for each fluorescent protein and the serum-PBS diluent were loaded in duplicate (technical replicates per standard) at 20 µL per well into black shallow 384-well plates (ProxiPlate-384 F Plus, 6008260, Revvity, Waltham, MA, USA), and fluorescence intensity of each standard and control was quantified using a PerkinElmer Victor Nivo^TM^ Multimode Plate Reader (Revvity) with the in-built 384-well plate settings. mCherry and tdTomato were measured at excitation 570/10 nm and emission 620/10 nm (measurement time 500 ms/well, flash energy 100 mJ, photomultiplier tube voltage 750 V). GFP was measured at excitation 480/30 nm and emission 530/30 nm (measurement time 500 ms/well, flash energy 10 mJ, photomultiplier tube voltage 488 V). All measurements were recorded as relative fluorescence units (RFUs). Please note that, although the 570/620 nm filter set was not optimal for tdTomato, these wavelengths were selected based on available instrumentation.

Standard curves were generated using four-parameter logistic regression (GraphPad Prism version 10). The lower limit of quantification (LLOQ) was defined as the lowest standard concentration meeting the following criteria: coefficient of variation (CV) <20% between replicate fluorescence measurements, signal-to-noise ratio (S/N) ≥10 and accuracy within 80-120%. The upper limit of quantification (ULOQ) was defined as the highest standard concentration with CV <15% before signal saturation. The dynamic range was determined as the concentration range between LLOQ and ULOQ (**Supplementary Fig. 3**).

Serum samples from experimental mice receiving AAV vector, plasmid DNA, or purified mCherry protein were diluted at 1:7.5 in PBS (1 part serum in 6.5 parts PBS) prior to fluorescence measurement. Naive sera collected from 5 untreated 8-week-old C57BL/6J male mice were diluted in PBS in the same manner and served as negative controls. Fluorescent protein concentrations in experimental samples and naive controls were quantified in triplicate (technical replicates per sample) by interpolation from standard curves generated with duplicate standards loaded on the same plate. Samples with fluorescence signals outside the dynamic range are reported as >ULOQ (exceeding the upper limit) or <LLOQ (below the lower limit). In our dataset, no samples exceeded the ULOQ.

### kidneyseg-pt

We devised kidneyseg_pt, a Python package for objective quantification of PT transduction using immunofluorescence microscopy images stained with LTL. Using RGB fluorescence microscopy images of kidney sections stained with LTL as inputs, kidneyseg_pt employs PT outer wall arc length-based partitioning with medial axis projection to quantify PT transduction percentage. The algorithm divides each PT area, defined as LTL-positive tubular structures, into a defined number (N) of partitions, classifies each partition as target (*e.g.*, fluorescent protein marker) signal-positive or negative, and determines the percentage of positive partitions, representing the PT transduction efficiency (**Fig. 7a, b**). In addition, kidneyseg_pt determines PT area (pixel^2^), total signal intensity (arbitrary unit, AU), average signal intensity per signal-positive pixel (AU/pixel), and average signal intensity in each PT area (AU/unit area or pixel^2^). We used N=50 in this study. Detailed kidneyseg_pt package workflow is provided in **Supplementary Methods**. The package is available on GitHub [https://github.com/nakaih-ohsu/AAV_Kidney_2026.git].

### Blinding

Experiments for Groups 6, 10, 13, 19, and 20, the marmoset study, and the injection in the first half of Group 1 mice were conducted in an unblinded fashion. All other studies were double-blinded. Although some experiments were conducted in an unblinded fashion, efforts were made to minimize bias and ensure objective observations.

### Statistics

Statistical analyses were performed using GraphPad Prism 10. Data are presented as mean ± standard deviation (SD) unless otherwise noted. To statistically demonstrate the presence of fluorescent protein in sera following agent administration, we compared fluorescent protein concentrations in sera between naive control sera (n = 5) and agent-injected mouse sera (n = 3 to 5 per group). The naive sample concentrations could be expressed as only “less than the lower limited of quantitation (LLOQ)”; therefore, the dataset did not satisfy assumptions of normality. Accordingly, we employed the two-tailed non-parametric Mann-Whitney U test to compare each agent-treated group and the naive control group. To account for multiple comparisons, p-values were corrected using the Benjamini-Hochberg False Discovery Rate (FDR) method. Similarly, the same non-parametric approach was applied to a statistical comparison between pseudotransduction and true transduction patterns. Statistical significance was defined as FDR-adjusted p-value (*i.e.*, q-value) of less than 0.05.

## Data availability

The raw and processed histology images generated in this study and have been deposited in Zenodo [https://doi.org/10.5281/zenodo.21431972]. The full uncropped western blot images are provided as **Supplementary Files 1-8**. The data generated and used in the figures are provided in the Source Data file.

## Code availability

The kidneyseg_pt Python package and sample images are available on GitHub [https://github.com/nakaih-ohsu/AAV_Kidney_2026.git]. The model weights for the kidneyseg_pt package, and images and masks used for kidneyseg_pt training, validation and analysis are available on Zenodo [https://doi.org/10.5281/zenodo.21431972].

## Acknowledgements

This work was supported in part by Public Health Service grants (R01DK146261 and R01DK144804), the American Society of Nephrology KidneyCure Bridge Grant Pilot Program, the Medical Research Foundation of Oregon Emergency Interim Support, and OHSU Presidential Bridge Funding to HN. The research reported in this publication used the computational infrastructure at the Oregon Health & Science University (OHSU) Advanced Computing Center (ARC), supported by the Office of Research Infrastructure Programs, Office of the Director, of the National Institutes of Health (NIH) under Award Number S10OD034224, and also used Bridges-2 at Pittsburgh Supercomputing Center through allocation BIO170079 from the Advanced Cyberinfrastructure Coordination Ecosystem: Services & Support (ACCESS) program, which is supported by National Science Foundation grants #2138259, #2138286, #2138307, #2137603, and #2138296. This marmoset work was supported by grants from the Program for Brain Mapping by Multidisciplinary Frontier Brain and Neuroscience Discoveries (Brain/MINDS 2.0; JP23wm0625001/JP24wm0625103 [to H.H.]) from the Japan Agency for Medical Research and Development (AMED); and by MEXT/JSPS KAKENHI (JP24K15743 [to Y.M.] and JP23H02791 [to H.H.]). We thank Guangping Gao and James M. Wilson for the AAV9 helper plasmid, Katja Pekrun and Mark A. Kay for the AAV-KP1 helper plasmid, Kei Adachi for assistance with manuscript preparation, Swapna Kollu for assistance with AAV vector production, Pratheppa Rajagopal for assistance with AAV vector genome quantification, Pretham Nandakumar for reviewing computer scripts, and the staff in the OHSU Advanced Light Microscopy shared resource for their expert technical assistance.

## Author contributions

A.S. and H.N. designed the study and wrote the manuscript. A.S. produced AAV vectors, processed mouse serum and tissue samples, performed fluorescence microscopy analysis, fluorometer measurements, and western blot analysis, and quantified vector genome copy numbers and vector genome transcripts. R.D., L.L., and H.N. coordinated and conducted C57BL/6J mouse experiments. R.D. also assisted in acquisition of microscopy images. A.S., T.G., M.H., and H.N. coordinated and conducted iMegKO mouse experiments. M.Y. and H.H. performed fluorescence microscopy analysis of marmoset samples. H.N. developed the kidneyseg_pt algorithm, wrote the kidneyseg_pt Python package, and performed computational analysis. All authors read and approved the final version of the manuscript.

## Competing interests

H.N. receives royalty of AAV-related technologies licensed by M34 Inc. H.N. serves as a consultant for biotech companies, is the founder of NeoSakura BTX, Inc., and holds shares of M34 Inc. NeoSakura BTX, Inc., and Sphere Gene Therapeutics. The remaining authors declare no competing interests.

## Supplementary Materials

Supplementary Materials (Supplementary Figures, Supplementary Methods, Supplementary Tables, Supplementary Data, Supplementary Discussion, and Supplementary Files), Source Data, and raw immunofluorescence microscopy image files and code deposited to Zenodo and GitHub, respectively, will be made publicly available upon publication of the peer-reviewed manuscript.

## References

1. Kohn, D. B., Chen, Y. Y. & Spencer, M. J. Successes and challenges in clinical gene therapy. Gene Ther 30, 738–746 (2023).

2. Khare, V. & Cherqui, S. Targeted gene therapy for rare genetic kidney diseases. Kidney Int 106, 1051–1061 (2024).

3. Peek, J. L. & Wilson, M. H. Cell and gene therapy for kidney disease. Nat Rev Nephrol 19, 451–462 (2023).

4. Davis, L. & Park, F. Gene therapy research for kidney diseases. Physiol Genomics 51, 449–461 (2019).

5. Rubin, J. D. & Barry, M. A. Improving Molecular Therapy in the Kidney. Mol Diagn Ther 24, 375–396 (2020).

6. Suarez-Amaran, L., Song, L., Tretiakova, A. P., Mikhail, S. A. & Samulski, R. J. AAV vector development, back to the future. Mol Ther 33, 1903–1936 (2025).

7. Wang, J. H., Gessler, D. J., Zhan, W., Gallagher, T. L. & Gao, G. Adeno-associated virus as a delivery vector for gene therapy of human diseases. Signal Transduct Target Ther 9, 78 (2024).

8. Liu, D., et al. Adeno-associated virus therapies: Pioneering solutions for human genetic diseases. Cytokine Growth Factor Rev 80, 109–120 (2024).

9. Lipkowitz, M. S., et al. Transduction of renal cells in vitro and in vivo by adeno-associated virus gene therapy vectors. J Am Soc Nephrol 10, 1908–1915 (1999).

10. Ikeda, Y., Sun, Z., Ru, X., Vandenberghe, L. H. & Humphreys, B. D. Efficient Gene Transfer to Kidney Mesenchymal Cells Using a Synthetic Adeno-Associated Viral Vector. J Am Soc Nephrol 29, 2287–2297 (2018).

11. Zincarelli, C., Soltys, S., Rengo, G. & Rabinowitz, J. E. Analysis of AAV serotypes 1-9 mediated gene expression and tropism in mice after systemic injection. Mol Ther 16, 1073–1080 (2008).

12. Furusho, T., et al. Enhancing gene transfer to renal tubules and podocytes by context-dependent selection of AAV capsids. Nat Commun 15, 10728 (2024).

13. Lang, J. F., Toulmin, S. A., Brida, K. L., Eisenlohr, L. C. & Davidson, B. L. Standard screening methods underreport AAV-mediated transduction and gene editing. Nat Commun 10, 3415 (2019).

14. Pekrun, K., et al. Using a barcoded AAV capsid library to select for clinically relevant gene therapy vectors. JCI Insight 4, e131610 (2019).

15. Hakui, H., Sairavi, A., Das, R., Furusho, T. & Nakai, H. Effective and Safe Gene Delivery to the Mouse Kidney via Slow Retrograde Renal Pelvis Injection of Adeno-Associated Virus Vectors. J Vis Exp 222, e67716 (2025).

16. Lin, M. H., Omachi, K., Begin, J. F., Richardson, J. L. & Miner, J. H. Mouse Alport podocytes are susceptible to AAV9 transduction in vivo. Kidney Int 109, 129–138 (2026).

17. Rocca, C. J., Ur, S. N., Harrison, F. & Cherqui, S. rAAV9 combined with renal vein injection is optimal for kidney-targeted gene delivery: conclusion of a comparative study. Gene Ther 21, 618–628 (2014).

18. Shen, X., et al. Transparenchymal Renal Pelvis Injection of Recombinant Adeno-Associated Virus Serotype 9 Vectors Is a Practical Approach for Gene Delivery in the Kidney. Hum Gene Ther Methods 29, 251–258 (2018).

19. Hillestad, M. L., Guenzel, A. J., Nath, K. A. & Barry, M. A. A vector-host system to fingerprint virus tropism. Hum Gene Ther 23, 1116–1126 (2012).

20. Konkalmatt, P. R., et al. Renal rescue of dopamine D2 receptor function reverses renal injury and high blood pressure. JCI Insight 1, e85888 (2016).

21. Chung, D. C., et al. Adeno-Associated Virus-Mediated Gene Transfer to Renal Tubule Cells via a Retrograde Ureteral Approach. Nephron Extra 1, 217–223 (2011).

22. Maturana, C. J. & Engel, E. A. Persistent transgene expression in peripheral tissues one year post intravenous and intramuscular administration of AAV vectors containing the alphaherpesvirus latency-associated promoter 2. Front Virol 4, 137991 (2024).

23. Picconi, J. L., et al. Kidney-specific expression of GFP by in-utero delivery of pseudotyped adeno-associated virus 9. Molecular therapy Methods & clinical development 1, 14014 (2014).

24. Mattar, C. N., et al. Systemic gene delivery following intravenous administration of AAV9 to fetal and neonatal mice and late-gestation nonhuman primates. FASEB J 29, 3876–3888 (2015).

25. Gonzalez, T. J., et al. Cross-species evolution of a highly potent AAV variant for therapeutic gene transfer and genome editing. Nat Commun 13, 5947 (2022).

26. Rosales, A., et al. Evolving adeno-associated viruses for gene transfer to the kidney via cross-species cycling of capsid libraries. Nat Biomed Eng 9, 1086–1100 (2025).

27. Peek, J. L., et al. Efficient kidney gene transfer and proximal tubule transduction using self-complementary AAV.cc47 vectors. Molecular therapy Methods & clinical development 33, 101606 (2025).

28. Garcia, F. G., et al. Integrated AAV optimization enables efficient gene delivery to kidney in murine and human tissue. Research Square, *Preprint, doi:* 10.21203/rs.3.rs-7925411/v1 (2025).

29. Peek, J. L., et al. Proximal Tubule-Targeted Adeno-Associated Virus (AAV) Gene Therapy for Cystinuria. J Am Soc Nephrol 35 (**10S**), (2024).

30. Alexander, I. E., Russell, D. W. & Miller, A. D. Transfer of contaminants in adeno-associated virus vector stocks can mimic transduction and lead to artifactual results. Hum Gene Ther 8, 1911–1920 (1997).

31. Seisenberger, G., et al. Real-time single-molecule imaging of the infection pathway of an adeno-associated virus. Science 294, 1929–1932 (2001).

32. Luni, C., Michielin, F., Barzon, L., Calabro, V. & Elvassore, N. Stochastic model-assisted development of efficient low-dose viral transduction in microfluidics. Biophys J 104, 934–942 (2013).

33. Madisen, L., et al. A robust and high-throughput Cre reporting and characterization system for the whole mouse brain. Nat Neurosci 13, 133–140 (2010).

34. Muzumdar, M. D., Tasic, B., Miyamichi, K., Li, L. & Luo, L. A global double-fluorescent Cre reporter mouse. Genesis 45, 593–605 (2007).

35. Chen, Z., et al. Transgenic zebrafish modeling low-molecular-weight proteinuria and lysosomal storage diseases. Kidney Int 97, 1150–1163 (2020).

36. Prutskova, N. P. & Seliverstova, E. V. Absorption capacity of renal proximal tubular cells studied by combined injections of YFP and GFP in Rana temporaria L. Comp Biochem Physiol A Mol Integr Physiol 166, 138–146 (2013).

37. Lacasse, V., Beaudoin, S., Jean, S. & Leyton, J. V. A Novel Proteomic Method Reveals NLS Tagging of T-DM1 Contravenes Classical Nuclear Transport in a Model of HER2-Positive Breast Cancer. Molecular therapy Methods & clinical development 19, 99–119 2020).

38. Wang, X., et al. Liver-secreted fluorescent blood plasma markers enable chronic imaging of the microcirculation. Cell Rep Methods 2, 100302 (2022).

39. Eshbach, M. L. & Weisz, O. A. Receptor-Mediated Endocytosis in the Proximal Tubule. Annu Rev Physiol 79, 425–448 (2017).

40. Cranfill, P. J., et al. Quantitative assessment of fluorescent proteins. Nat Methods 13, 557–562 (2016).

41. Katayama, H., Yamamoto, A., Mizushima, N., Yoshimori, T. & Miyawaki, A. GFP-like proteins stably accumulate in lysosomes. Cell Struct Funct 33, 1–12 (2008).

42. Liu, F., Song, Y. & Liu, D. Hydrodynamics-based transfection in animals by systemic administration of plasmid DNA. Gene Ther 6, 1258–1266 (1999).

43. Fumoto, S., Nishimura, K., Nishida, K. & Kawakami, S. Three-Dimensional Imaging of the Intracellular Fate of Plasmid DNA and Transgene Expression: ZsGreen1 and Tissue Clearing Method CUBIC Are an Optimal Combination for Multicolor Deep Imaging in Murine Tissues. PLoS One 11, e0148233 (2016).

44. Woodard, L. E., et al. Hydrodynamic Renal Pelvis Injection for Non-viral Expression of Proteins in the Kidney. J Vis Exp, 56324 (2018).

45. Kobayashi, N., Nishikawa, M., Hirata, K. & Takakura, Y. Hydrodynamics-based procedure involves transient hyperpermeability in the hepatic cellular membrane: implication of a nonspecific process in efficient intracellular gene delivery. J Gene Med 6, 584–592 (2004).

46. Wang, M. Cross-species evolution of capsid libraries for gene delivery to the kidneys. Nat Rev Nephrol 21, 217 (2025).

47. Orabi, D., Sharma, D. & Gardner, J. M. Cross-species evolution of kidney-targeting adeno-associated virus vectors for ex vivo organ editing. Am J Transplant 26, 1209–1210 (2026).

48. Hickman, P. G., Asokan, A. & Wilson, M. H. Rewriting Renal Fate: The Evolving Landscape of Adeno-Associated Virus-Mediated Kidney Gene Therapies. Kidne*y360* 7, 422–425 (2026).

49. Christensen, E. I., Birn, H., Storm, T., Weyer, K. & Nielsen, R. Endocytic receptors in the renal proximal tubule. Physiology (Bethesda) 27, 223–236 (2012).

50. Nielsen, R., Christensen, E. I. & Birn, H. Megalin and cubilin in proximal tubule protein reabsorption: from experimental models to human disease. Kidney Int 89, 58–67 (2016).

51. Mori, K. P., et al. Increase of Total Nephron Albumin Filtration and Reabsorption in Diabetic Nephropathy. J Am Soc Nephrol 28, 278–289 (2017).

52. Matsushita, K., et al. Cilastatin Ameliorates Rhabdomyolysis-induced AKI in Mice. J Am Soc Nephrol 32, 2579–2594 (2021).

53. Ulrix, W., Swinnen, J. V., Heyns, W. & Verhoeven, G. The differentiation-related gene 1, Drg1, is markedly upregulated by androgens in LNCaP prostatic adenocarcinoma cells. FEBS Lett 455, 23–26 (1999).

54. Chen, S., et al. Multi-omic and spatial analysis of mouse kidneys highlights sex-specific differences in gene regulation across the lifespan. Nat Genet 57, 1213–1227 (2025).

55. Matsuzaki, Y., et al. Systematic evaluation of BBB-penetrant AAV capsids in marmosets identifies VCAP-102 as a highly efficient brain-transducing capsid. Mol Ther Adv 34, 201783 (2026).

56. Goto, S., Hosojima, M., Kabasawa, H. & Saito, A. The endocytosis receptor megalin: From bench to bedside. Int J Biochem Cell Biol 157, 106393 (2023).

57. Shaner, N. C., Patterson, G. H. & Davidson, M. W. Advances in fluorescent protein technology. J Cell Sci 120, 4247–4260 (2007).

58. Balazsi, G., van Oudenaarden, A. & Collins, J. J. Cellular decision making and biological noise: from microbes to mammals. Cell 144, 910–925 (2011).

59. Hahl, S. K. & Kremling, A. A Comparison of Deterministic and Stochastic Modeling Approaches for Biochemical Reaction Systems: On Fixed Points, Means, and Modes. Front Genet 7, 157 (2016).

60. Jackson, B. & Oken, D. E. Internephron heterogeneity of filtration fraction and disparity between protein-and hematocrit-derived values. Kidney Int 21, 309–315 (1982).

61. Frommer, J. P., Laski, M. E., Wesson, D. E. & Kurtzman, N. A. Internephron heterogeneity for carbonic anhydrase-independent bicarbonate reabsorption in the rat. J Clin Invest 73, 1034–1045 (1984).

62. Caplanusi, A., et al. Intravital multi-photon microscopy reveals several levels of heterogeneity in endocytic uptake by mouse renal proximal tubules. J Cell Mol Med 12, 351–354 (2008).

63. Ghaffarzadeh-Esfahani, M. & Gheisari, Y. Capsid Engineering of Adeno-Associated Viruses for Targeted Gene Therapy in Kidney Diseases. Hum Gene Ther 37, 819–832 (2026).

64. Asokan, A. & Wilson, M. H. Filtering through AAV capsid libraries for effective kidney gene transfer. Kidney Int 107, 591–593 (2025).

65. Viecelli, H. M., et al. Treatment of phenylketonuria using minicircle-based naked-DNA gene transfer to murine liver. Hepatology 60, 1035–1043 (2014).

66. Earley, L. F., et al. Adeno-associated Virus (AAV) Assembly-Activating Protein Is Not an Essential Requirement for Capsid Assembly of AAV Serotypes 4, 5, and 11. J Virol 91, e01980–019816 (2017).

67. Powers, J. M., Chang, X. L., Song, Z. & Nakai, H. A Quantitative Dot Blot Assay for AAV Titration and Its Use for Functional Assessment of the Adeno-associated Virus Assembly-activating Proteins. J Vis Exp 136, 56766 (2018).

68. Adachi, K., Enoki, T., Kawano, Y., Veraz, M. & Nakai, H. Drawing a high-resolution functional map of adeno-associated virus capsid by massively parallel sequencing. Nat Commun 5, 3075 (2014).

